# Heritable and sex-specific variation in the development of social behavior in a wild primate

**DOI:** 10.1101/2022.10.21.513189

**Authors:** Elizabeth C. Lange, Madison Griffin, Arielle S. Fogel, Elizabeth A. Archie, Jenny Tung, Susan C. Alberts

**Author notes:** Author Contributions:* ECL: conceptualization, methodology, formal analysis, writing (original draft & review/editing). MG: formal analysis, writing (review/editing). ASF: formal analysis, writing (review/editing). JT: resources, data curation, writing (review/editing), funding acquisition. EAA: resources, data curation, writing (review/editing), funding acquisition. SCA: conceptualization, supervision, resources, data curation, writing (review/editing), funding acquisition.

## Abstract

Affiliative social bonds are linked to fitness components in many social mammals. However, despite their importance, little is known about how the tendency to form social bonds develops in young animals, or if the development of social behavior is heritable and thus can evolve. Using four decades of longitudinal observational data from a wild baboon population, we assessed the environmental determinants of an important social developmental milestone in baboons—the age at which a young animal first grooms a conspecific—and we assessed how mother-offspring grooming reciprocity develops during the juvenile period. We found that grooming development differs between the sexes: female infants groom at an earlier age and reach reciprocity in grooming with their mother earlier than males. Using the quantitative genetic ‘animal model’, we also found that age at first grooming behavior for both sexes is weakly heritable (h^2^ = 4.3%). These results show that sex differences in grooming emerge at a young age; that strong, reciprocal social relationships between mothers and daughters begin very early in life; and that age at first grooming is heritable and therefore can be shaped by natural selection.

## Background

In humans and other mammals, individuals with more numerous or stronger social bonds in adulthood are often healthier and live longer, suggesting that strong social relationships should be favored via natural selection (1, 2). Social relationships in early life are also important, as they represent crucial opportunities to develop social skills and may have long-lasting consequences in their own right. For instance, in many species, positive social environments early in life are linked to better social relationships in adulthood (3–7), better health (3, 8), and increased longevity (9–13). In primates, social bonds are commonly developed and maintained through social grooming, a primary affiliative behavior in many species, including baboons (14–17). Furthermore, stronger grooming relationships and/or a higher frequency of grooming have been linked to longer lifespans in both male and female baboons (14, 18–20), suggesting that the development of grooming behavior is important to fitness in this species. Investigating the development of grooming behavior can therefore shed light on how environmental and genetic sources of variance contribute to the emergence of a social behavior with clear links to fitness.

Sex differences are common in the development of social behavior – including social grooming – and are pronounced in primates (21–23). In several species of non-human primates, juvenile females invest more in grooming relationships, while juvenile males invest more in play and agonistic behavior (22, 24–26). Such sex differences may reflect differences in future fitness benefits for each sex. For example, in species such as yellow baboons and rhesus macaques, in which females remain in their natal group and males disperse, the grooming partners of immature females will often be potential long-term partners, while the same is not true of males. In addition, in many such species, females experience nepotistic rank inheritance, which involves support from family members, while males must acquire adult dominance rank via physical competition without familial support (27–29). Thus, behaviors that promote future fighting ability may be more essential during development for males than for females (21–23).

Because social grooming is linked to fitness, it should evolve by natural selection if variation in grooming behavior has a genetic component. The genetic basis of trait variation is often assessed by measuring a trait’s narrow-sense heritability (h^2^), the proportion of phenotypic variation explained by additive genetic effects (30). In humans, many traits related to social interactions have a heritable component (31–34). In non-human vertebrates, research is more limited but generally supports the idea that variation in social behavior has measurable heritability (35–44). However, no study has assessed whether the onset and development of social behavior is heritable in wild populations. Therefore, the relative contributions of genes versus the environment, and the degree to which developmental features of social behavior can be targeted by natural selection, remain unclear.

Here, we examined the development of grooming behavior in infants living in the well-studied baboon population of the Amboseli basin (45). We had four goals. First, we examined the socio-environmental determinants of the age at which a young animal first groomed a conspecific. Second, we determined the identity of the recipient of each subject’s first grooming efforts. Because infants are in frequent proximity to their mothers, who are important providers of nutrition and information, we predicted that subjects would disproportionately choose their mothers as the recipients of their first grooming efforts. Third, to determine how grooming relationships develop, we examined how grooming reciprocity with mothers changed across the juvenile period and into early adulthood. Because young animals lack the coordination and skill to perform grooming behaviors, we predicted that when offspring are young, mothers would initiate grooming more, but that the mother-offspring grooming relationship would become more equitable during development. We also hypothesized that mother-daughter relationships would be more reciprocal than mother-son relationships because in the Amboseli baboons, as in many primates, mothers are an important social partner to females in adulthood (15, 23, 46). In contrast, sons disperse from their natal social group as they approach maturity, terminating mother-son relationships. Finally, we measured the heritability of age at first grooming, using the ‘animal model,’ a linear mixed effects model that estimates both environmental and genetic sources of variance and covariance in phenotypes (47, 48).

## Methods

### Study population and study subjects

The Amboseli Baboon Research Project (ABRP) – a longitudinal study of wild yellow and yellow-anubis admixed baboons (*Papio cynocephalus × P. anubis*) living in and around Amboseli National Park, Kenya – has collected behavioral, environmental, and demographic data from individually recognized baboons on a near-daily basis since 1971. The ABRP has also collected tissue samples since 1989 and fecal samples for DNA analysis since 2000 (45). Baboons in Amboseli live in stable social groups containing multiple adults and juveniles of both sexes, ranging in size from approximately 20 to 100 animals. The ABRP monitors multiple such groups (‘study groups’) in the Amboseli ecosystem.

Our study subjects included 781 immature baboons (368 males and 413 females) born into Amboseli study groups between 1983 and 2020. Ages of all subjects were known to within a few days’ error. We included the few subjects whose mothers died before the subject began to groom conspecifics (n=11 of 781 subjects). We excluded individuals who died before one year of age and those whose first grooming was known or likely to have occurred during a period of reduced data collection or during group fissions (i.e., animals between 0.58 - 2.6 years of age during these periods, the timespan during which most first grooming events occur).

Study subjects were habituated to experienced observers who recognize individual baboons by sight. Age at first grooming was defined as a subject’s age in years at the earliest interaction in which it was observed to perform coordinated and systematic picking through the fur of another animal. Data were collecting using an approach that we refer to as “representative interaction sampling,” which is designed to avoid biases from uneven sampling of study subjects. Specifically, an observer moves systematically through the group while carrying out 10-minute focal animal samples according to a predefined, randomized list of focal animals. They simultaneously record all grooming interactions in their line of sight, whether or not they involve the focal animal. Importantly, while this approach is designed to mitigate bias, it cannot capture all grooming interactions (e.g., grooming may occur outside of an observer’s line of sight, or when observers are not watching the group). Consequently, the age at which we first observed a subject to groom (our measurement of age at first grooming) should be considered the latest date by which this milestone was achieved, rather than the exact date on which it was achieved. This ascertainment error is unlikely to be correlated with other predictors in our models except for group size, which affects the ratio of animals to observers in a group and hence per-animal observation effort. Because we are less likely to observe a given grooming interaction in a large group compared to a small group, we included a measure of observer effort in our models of age at first grooming below.

### Environmental predictors of age at first grooming

We assessed a range of maternal, social, and physical environmental variables as potential predictors of age at first grooming, chosen because they are known or likely contributors to infant development, described in detail in Supplementary Methods and Analyses (see “*Description of predictors used in our model selection analysis*”). These included the subject’s own sex (male or female) as well as: (i) maternal parity, (ii) maternal social isolation, (iii) maternal proportional dominance rank, (iv) maternal alpha rank status (whether or not the subject’s mother was the highest-ranking female in the group at the subject’s birth), (v) season of birth (wet or dry), (vi) presence of drought in the first year of life, (vii) group size at birth, and (viii) presence of maternal siblings between birth and first grooming. In addition, because the Amboseli baboons are an admixed population (49, 50), we conducted secondary analyses that included genetic hybrid scores as fixed effects in our models. Because these secondary analyses included a smaller set of individuals (i.e., those for whom these scores are available) and produced similar results to our main models, we present them in the Supplementary Methods and Analyses (see *“Effects of hybrid score on age at first grooming behavior*”).

### Modeling environmental predictors of age at first grooming

#### LMM to examine sex differences in age at first grooming

Visual inspection of the data indicated that females started grooming earlier than males, so we first assessed the effects of sex on age at first grooming using a linear mixed model. We included random effects of social group ID, maternal ID, and year of birth, using the Amboseli ‘hydrological year’, which begins in November with the annual rainy season and continues through October of the following year (e.g., hydrological year 2021 began in November 2020 and continued through October 2021). We fit this model using the ‘glmmTMB’ R package (51).

#### Model selection to examine maternal, social, and physical environmental predictors of age at first grooming

To analyze the larger set of potential predictors of age at first grooming, we used a model selection approach that compared a set of mixed effects Cox proportional hazard models that included different combinations of these variables. We modeled age at first grooming as time-to-event data that were right-censored, with subject’s age as one of the predictors. We examined the predictors of age at first grooming separately for males and females because preliminary analyses showed non-proportional hazards of age at first grooming when both sexes were combined in the same model.

We used a model selection approach because we did not have specific predictions about which of our predictors would best explain variation in age at first grooming (52, 53). Specifically, we used the “coxme” function from the R coxme package (54) to evaluate a set of candidate mixed effects Cox proportional hazards models that included linear fixed effects of maternal social isolation, maternal parity, maternal proportional rank, maternal alpha status, drought, season of birth, group size, and presence of maternal sibling. We also included a fixed effect of observer effort, which we calculated as the number of grooming events collected per group member per month for each group, averaged over the time period from birth to first grooming for each subject (see “*Study population and study subjects*”). These models also included random effects of social group, hydrological year of birth, and maternal ID.

To compare models, we calculated adjusted Akaike’s Information Criterion (AIC_c_, 55) for combinations of fixed effects using the “dredge” function from the MuMIn R package (56). We calculated model-averaged parameter estimates for fixed effects using only ‘top models’ where ΔAIC_c_ values were within two units of the best model with the lowest AIC_c_ value (52, 53). Estimates were calculated from the full coefficient set using the “model.avg” function from the MuMIn package, where terms that were not included in a model were set to zero.

### Recipients of first observed grooming effort (‘first grooming partner’)

We identified the recipients of all 781 subjects’ first grooming efforts (i.e., their ‘first grooming partners’). First, we used a binomial test to test the prediction that mothers were disproportionately the recipients of subjects’ first grooming efforts, compared to all other social group members. Second, we used a Chi square test to test for a sex difference in the likelihood of grooming the mother compared to any other social group member. Here, we compared the first grooming partners of males (grouped into mothers versus non-mother females and males) and the first grooming partners of females (similarly grouped). Third, we tested for sex differences in grooming the mother compared to non-mother females. Here, we compared the first grooming partners of male subjects (grouped into mothers and non-mother females) to female subjects (similarly grouped), again using a chi square test. Finally, we tested the prediction that relatives (r ≥ 0.0625; see below) were disproportionally the subject’s first grooming partner using a binomial test.

We categorized first grooming partners by relatedness using the “kinship” function in the kinship2 R package (57), which estimates relatedness coefficients between dyads based on multi-generation pedigree information (our full pedigree includes 1866 individuals; 41, 58). All 781 of our subjects had known mothers and 489 had known fathers. The number of fathers and paternal kin identified as first grooming partners is underestimated because 292 subjects lacked paternity assignments.

### Mother-offspring grooming reciprocity

We defined grooming reciprocity as the number of grooming events initiated by the mother divided by the total number of grooming events between each mother-offspring dyad, assessed monthly during the juvenile period. Higher values of this metric correspond to cases where the mother initiated more grooming, while lower values correspond to cases where the offspring initiated more grooming; values of 0.5 represent perfect reciprocity. Our dataset excluded months during several periods of reduced data collection or during group fissions.

Females and males reach full maturation at different ages: females typically reach menarche – which we consider the onset of adulthood for female baboons – in their fifth year of life (59). In contrast, while males typically reach puberty (testicular enlargement) in their sixth year of life, they do not attain full adult dominance rank and begin mating until their eighth year of life. To account for these developmental differences, we split our analyses into two time periods; first, we assessed mother-offspring grooming reciprocity from birth to the median age of female menarche (4.5 years), a time period when both males and females were immature, and second, we assessed mother-offspring grooming reciprocity from the median age of menarche to median age of male rank attainment (7.7 years), a time period when females were fully adult but most males were not.

To assess reciprocity, we used binomial mixed effects models with a logit link using the “glmmTMB” function in the glmmTMB R package. We assessed fixed effects of age (both linear and squared), offspring sex, and their interaction. To control for repeated measures and maternal effects, we included offspring ID nested within maternal ID as a random effect. Social group and hydrological year of birth were also modelled as random effects. Our dependent variable was the number of mother-initiated grooming events divided by the number of total mother offspring grooming events in a given month for each mother-offspring pair. To assess which partner drove changes in mother-offspring reciprocity over time, we built separate post hoc models of the number of grooming events initiated by the mother, the number of grooming events initiated by the offspring, and the total number of grooming events between each mother-offspring dyad in mixed effects models with a zero inflated negative binomial distribution using the same fixed and random effects in the reciprocity model. We confirmed our results were robust to the timing of dispersal by adult male offspring, in case males with stronger maternal bonds delayed dispersal (see Supplementary Methods and Analyses).

### Genetic variance in age at first grooming

To measure the heritability of age at first grooming, we combined males and females together and used the ‘animal model,’ a linear mixed effects model that combines pedigree information with phenotypic values (47, 48). In the ‘animal model’, a vector of individual phenotypes is the response variable with estimates of each individual’s breeding value included as a random effect. The covariance of breeding values among individuals is, in turn, affected by trait heritability and relatedness in the sample. Here, we derived relatedness values from the multigenerational Amboseli pedigree, subset to 1315 individuals and a maximum of 6 generations (i.e., the subset of the full pedigree necessary to estimate *r* for animals in this data set). Among these 1315 individuals, 871 had known mothers and 531 had known fathers. Our dataset included 97 full sibling pairs, 905 maternal half sibling pairs, 1367 paternal half sibling pairs, and 613 sibling pairs that were at least maternal half siblings but for whom the father of at least one of the pair was unknown (i.e., some of these pairs could have been full siblings).

In the animal model, environmental effects are important to include as fixed effects because these effects can influence heritability estimates. We ran two separate animal models with fixed effects, using our model selection of predictors of age at first grooming to inform our decisions about which fixed effects to include in these animal models. In our first model (the ‘full model’) we included as fixed effects all variables that appeared in more than half of the top models for either males or females: subject’s sex, social group size, observer effort, presence of siblings, drought by sex interaction, maternal alpha rank by sex interaction, and maternal proportional rank by sex interaction. We then ran a second ‘reduced model’ that included sex, social group size, and observer effort as fixed effects because these three parameters were included in all top models in the environmental determinants analysis in both sexes (see Results).

For both the full and reduced models, we modeled a nested set of random effects, a standard procedure for validating heritability estimates with the animal model: (1) a ‘maternal effects’ only model that included only the random effect of maternal ID, (2) an ‘early environmental effects’ model that included the random effects of maternal ID, social group, and hydrological year of birth to account for cohort effects, and (3) a ‘heritability’ model that included a random effect of animal breeding value along with maternal ID, social group, and cohort effects.

Including environmental effects as fixed effects in an animal model can reduce residual variance and thus alter heritability estimates (60). Therefore, we also report an ‘ intercept only’ model that did not include fixed effects but did include all four random effects. In addition, in our calculations of heritability from models with fixed effects, we also included the variance explained by fixed effects in our estimates of phenotypic variance (61, 62).

Because the subject’s genetic hybrid score was a potential predictor of age at first grooming (see Results), we also ran a secondary set of animal models that included the subject’s genetic hybrid score in addition to environmental predictors. However, the number of subjects with hybrid scores was small (N=237 versus N=781 for the full data set), limiting our power for this analysis (see Supplementary Methods and Analyses).

We implemented the animal model using the MCMCglmm R package (63). We used weakly informative priors (V=1, nu=0.002) and a total of 30,000,000 iterations with a burn-in period of 500,000 and a thinning of 10,000. Our effective sample sizes for fixed effects exceeded 2,590 and those for random effects exceeded 2,050 in all models. We compared model fit using Deviance Information Criteria (DIC) scores, which is interpreted similarly to an AIC score (64, 65). Because variance component estimates are constrained to be positive, we used DIC comparisons of models with and without the random effect of animal to assess if heritability was significantly different than zero.

## Results

### Both sex and environmental variables predicted age at first grooming

Females were observed to groom earlier than males (Figure S1; GLMM: b=0.189±0.025, z=0.760; p<0.001), with females reaching age at first grooming at an average age of 0.9 years (10.7 months; 95%CI: 10-11.5 months) and males reaching age at first grooming at an average age of 1.1 years (13 months; 95%CI: 12.2-13.8 months). Our model selection using Cox proportional hazards models for females and males separately yielded 11 models with ΔAIC_c_ values <2 from the model with the lowest AIC_c_ value; we refer to these at the ‘top models’ (Table S1).

Social group size appeared as a fixed effect in all top models for both sexes (Table S1), with males and females in smaller groups exhibiting earlier age at first grooming (Figure S2). Females in the smallest groups groomed a median of 0.2 years (2.4 months) earlier than those in the largest groups (HR=0.989, 95%CI: 0.982-0.996; Table S2a, Figure S2a). Males in the smallest groups groomed a median of 0.3 years (4 months) earlier than those in the largest groups (HR=0.985, 95%CI: 0.975-0.995; Table S2b, Figure S2b).

For females, drought in the first year of life also appeared in all top models (Table S1a): females who experienced drought in the first year of life groomed earlier than those who did not (HR: 1.498, 95%CI: 1.041-2.155; Table S2a; Figure S3). However, the median age at first grooming for females who experienced drought (0.91 years) was almost identical to those who did not experience drought (0.89 years), suggesting a small effect of drought that may primarily affect subjects who first groomed after the median age (Figure S3). Maternal social isolation, presence of maternal sibling, maternal parity, season of birth, and maternal proportional rank each appeared in only one of the top models for females, and model-averaged parameters showed hazard ratios that overlapped one, suggesting no significant effect of these variables (Tables S1a, S2a).

For males, maternal proportional rank and maternal alpha status were included in 4 of 5 top models (Table S1b). Males with alpha mothers were 1.5 times more likely to achieve first grooming at any age compared to males with non-alpha mothers (HR: 1.473, 95%CI: 1.048-2.410; Table S2b; Figure S4b). However, controlling for alpha status, males with high-ranking mothers were *less* likely to achieve first grooming at any age compared to males with low-ranking mothers (HR: 0.684, 95%CI: 0.416-0.970; Table S2b; Figure S4a). Drought in the first year of life, presence of a maternal sibling, and maternal parity were each included in only one top model for males, but model-averaged parameters for these effects showed hazard ratios that overlapped one (i.e., no effect; Tables S1b, S2b).

Observer effort – a technical rather than biological variable – appeared in all the top models for both sexes in the direction we expected: individuals were observed grooming at earlier ages when observer effort was greater (females: HR: 1.175, 95%CI: 1.099-1.256; males: HR: 1.132, 95%CI: 1.049-1.220; Tables S1, S2; Figure S5).

Our secondary analyses including the mother’s hybrid score or the subject’s hybrid score yielded weak evidence that the subject’s hybrid score, but not the mother’s, contributed to variation in age at first grooming (Supplementary Methods and Analyses, Tables S3-S6). Specifically, subject’s hybrid score appeared in approximately half the top models for both sexes (Table S3), but the confidence intervals for the model-averaged hazard ratio for subject’s hybrid score were very large and overlapped one for both sexes (Table S4; females: HR: 7.648, 95%CI: 0.680-85.983; males: HR: 7.716, 95%CI: 0.633-95.097).

### Recipients of first observed grooming effort (‘first grooming partner’)

First grooming partners were disproportionately females: 67% of female subjects (n=277) and 69% of male subjects (n=255) were first observed grooming a female, while 33% of female subjects (n=113) and 31% of male subjects were first observed grooming a male (n=136; Figure 1). Mothers were by far the most common first grooming partner for both males and females, and both sexes chose to groom their mother first significantly more often than expected by chance (Figure 1a; observed=0.328, expected=0.017, p<0.0001). However, a clear sex difference emerged in which male subjects groomed their mothers first more frequently than female subjects (of all potential partners, 37% of males’ first partners were their mother vs 29% for female subjects; Figure 1a; χ^2^=5.16, d.f.=1, p=0.023). At the same time, non-mother females were more common first partners for females than for males (38% of first partners for females vs 32% of first partners for males were non-mother females; Figure 1a; *χ*^2^=4.94, d.f.=1, p=0.026).

In the aggregate, 59% of subjects were first observed grooming kin with r ≥ 0.0625, a significantly greater percentage than expected by chance (Figure S6; expected=0.259, p<0.0001); 48% of subjects chose close kin (r≥0.25). While males made up a smaller proportion of first grooming partners than females overall (Figure 1b), the proportion of close kin was similar among male and female first grooming partners. However, in spite of the importance of close kin in general, fathers were rarely first grooming partners: they represented a minimum of ~1% of first grooming partners (n=13, Figure 1b) and a maximum of ~6% of first grooming partners (n=49, if all first grooming partners in the ‘unknown adult male’ category were fathers).

**Figure 1.**
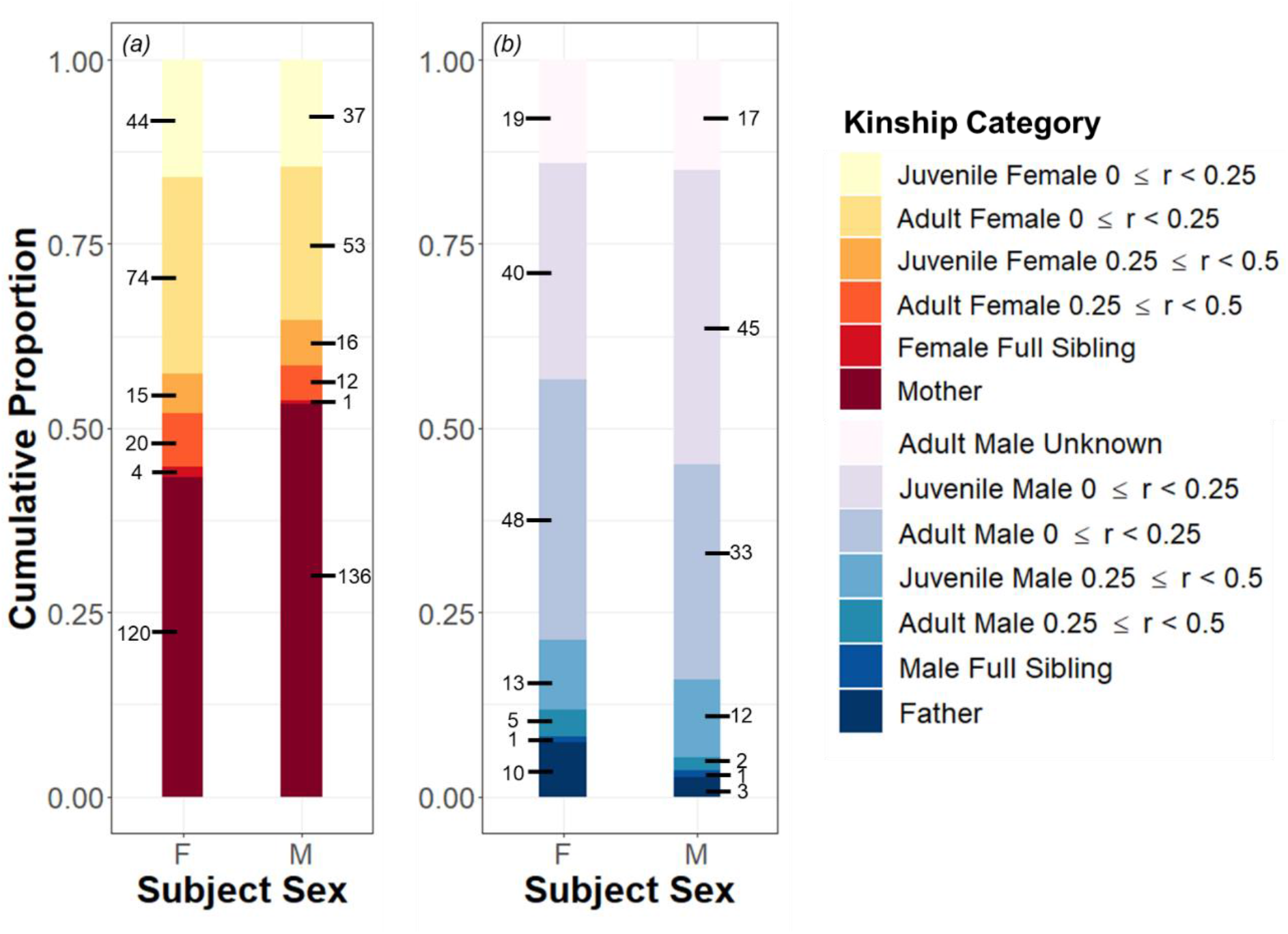
First grooming partners for study subjects. Numbers next to each bar denote the number of first grooming partners in that category. (a) First grooming partners who were female. (b) First grooming partners who were male. Age categories for first grooming partners included (i) juvenile males (aged <7 years, includes subadults), (ii) juvenile females (age <4 years), (iii) adult males, and (iv) adult females. Kinship categories included r=0.5 (e.g., parents, full siblings), 0.25 ≤ r < 0.5 (e.g., half siblings, grandparents, full aunts and uncles), r<0.25 (e.g., cousins, half aunts and uncles, half nieces and nephews, more distantly related kin, and unrelated partners), and unknown (applies only to immigrant males with no known offspring or adult relative in the group). The number of fathers and paternal relatives as first grooming partners is underestimated, because 292 subjects lacked paternity assignments. More fine-grained kinship categories are presented in Figure S6.

### Mother-offspring grooming reciprocity

Mother-daughter dyads were significantly more reciprocal than mother-son dyads overall (Figure 2, Table 1). Furthermore, daughters reached full reciprocity with mothers at 3.2 years, while males did not reach full reciprocity with mothers until 5 years of age (Figure 2). Relationships between mothers and daughters were more reciprocal than mother-son relationships during both the juvenile period for both sexes (birth – 4.5 years) and the period corresponding to subadulthood in males (4.5 – 7.7 years), but the sexes converged in mother-offspring reciprocity by 8 years of age (Figure 2).

**Figure 2.**
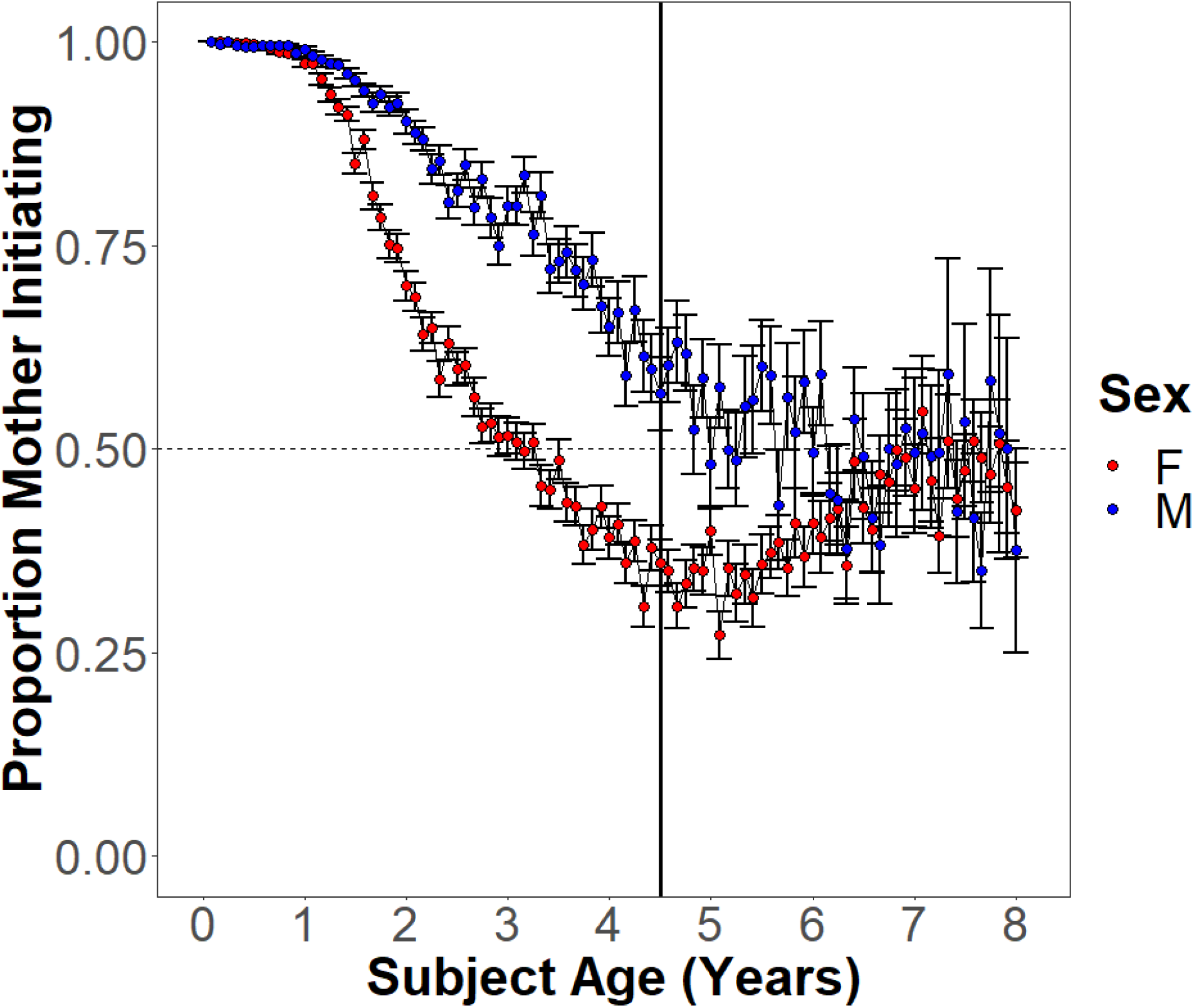
Proportion of mother-offspring grooming events where the mother-initiated grooming events for sons (blue) and daughters (red). The dashed horizontal line indicates perfect reciprocity where the number of grooms by the mother and offspring are equal. The solid vertical black line is at 4.5 years (median age at female menarche). Points are monthly averages and bars denote standard error.

**Table 1.** Predictors of mother-offspring grooming reciprocity: results of binomial generalized linear mixed models for two different time periods. Age (in months) was centered and standardized before analyses so estimates represent change in grooming reciprocity for one SD increase in age. Significant effects are bolded.

| Age Range | Effect | estimate | SE | Z value | P value | Interpretation |
| --- | --- | --- | --- | --- | --- | --- |
| a) 0-4.5 years | **Sex (male)** | **0.247** | **0.015** | **16.47** | **<0.001** | Females are more reciprocal with mothers than are males |
|  | **Age** | **-0.351** | **0.007** | **-47.02** | **<0.001** | Older animals are more reciprocal than younger animals |
|  | **Age<sup>2</sup>** | **-0.031** | **0.008** | **-3.81** | **<0.001** | Reciprocity is approached more rapidly with increasing age |
|  | **Sex\*Age** | **0.187** | **0.012** | **15.70** | **<0.001** | Reciprocity is achieved more rapidly in females than males |
|  | Sex*Age<sup>2</sup> | -0.015 | 0.012 | -1.25 | 0.213 |  |
| b) 4.5-7.7 years | **Sex (male)** | **0.162** | **0.059** | **2.75** | **0.006** | Males are more reciprocal with mothers than are females (i.e., females disproportionately initiate grooming with mothers) |
|  | **Age** | **0.101** | **0.020** | **5.13** | **<0.001** | Older animals are more reciprocal than younger animals |
|  | Age<sup>2</sup> | 0.006 | 0.022 | 0.27 | 0.785 |  |
|  | **Sex\*Age** | **-0.160** | **0.041** | **-3.86** | **<0.001** | With increasing age, females return to reciprocity with mothers after disproportionately being initiators; males maintain reciprocity with mothers during this period |
|  | Sex*Age<sup>2</sup> | 0.026 | 0.044 | 0.60 | 0.548 |  |

Sex differences in mother-offspring grooming relationships were primarily driven by sex differences in the offspring’s behavior rather than the mother’s behavior (Figure S7, Table S7). During infancy, mothers groomed male and female offspring at similar relative frequencies, and although mothers groomed female offspring at a slightly higher frequency than male offspring starting around 2.1 years of age, these differences remained relatively small throughout the period of maturation (Figure S7a). In contrast, despite the fact that males were more likely to target their mothers for first grooming than females, female offspring directed grooming towards their mothers at a much greater relative frequency than male offspring. This pattern started early in life and persisted throughout the maturation period (Figure S7b).

### Genetic variance in age at first grooming

Heritability of age at first grooming was 0.043 in the best fitting models (95% credible interval: 0.002-0.110), which included heritability along with fixed effects (the reduced model and full model fit equally well; Figure 3; Tables 2, S8). Nested models that did not include genetic effects (i.e., the ‘maternal effects’ and ‘environmental effects’ models in Tables 2, S8) had ΔDIC > 2 from models with additive genetic effects, suggesting that while heritability is low, models including genetic effects significantly improve fit over ones without them (Tables 2, S8). In the model with no fixed effects, the heritability estimate was nearly identical to the model with fixed effects, and the confidence intervals for heritability in the two models was very similar (h^2^=0.047, 95%credible interval: 0.002-0.122; Table 2), suggesting that the inclusion of fixed effects did not confound heritability estimates. In the model that included the subset of individuals with hybrid scores, the point estimate of heritability was higher (h^2^=0.125, 95% credible interval: 0.001-0.307; Table S9; Figure S8), but the credible intervals of heritability estimates were larger and overlapped completely with those in the model without hybrid scores.

**Figure 3.**
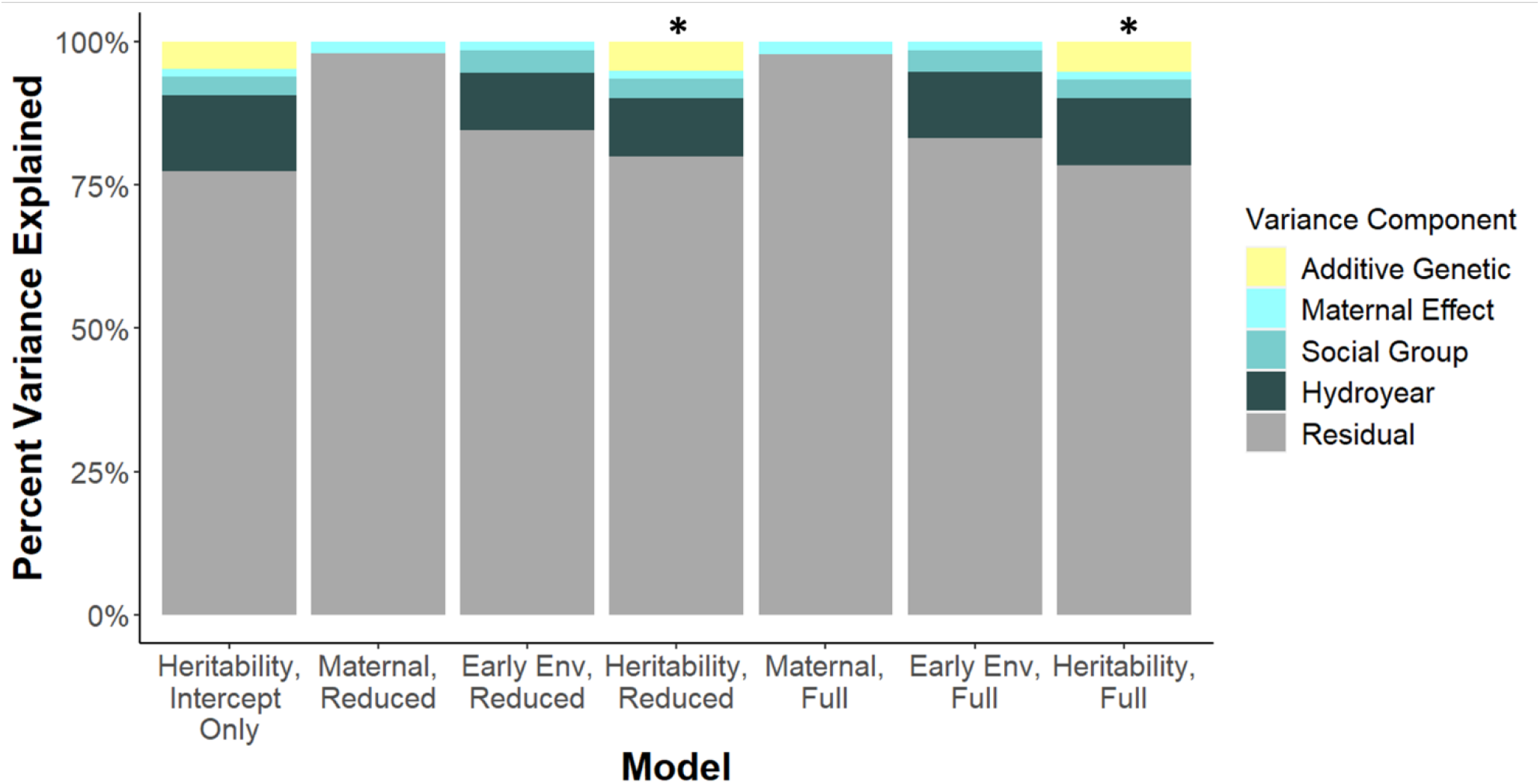
Percent of variance in age at first groom explained by additive genetic, maternal, social group, hydrological year, and residual effects. The best fitting models (indicated by a star) included additive genetic effects, maternal effects, social group effects, hydrological year effects as well as fixed effects. All models with additive genetic effects fit better than nested models that did not model additive genetic effects. See Tables 2, S8, and methods for definitions of models.

**Table 2.** Comparison of nested animal models. (i) Intercept-only model: only an intercept is included as a fixed effect. (ii) Maternal effects, reduced model: only maternal ID (m) was included as a random effect, with sex (s), social group size (gs), and observer effort (oe) as fixed effects. (iii) Environmental effects models, reduced model: maternal ID (m), social group ID (g), and cohort (c, based on hydrological year of birth) were included random effects. Fixed effects are as in (ii). (iv) Heritability, reduced model: animal ID (a, to estimate the breeding value) was included as a random effect, in addition to maternal ID, social group ID, and cohort. Fixed effects are as in (ii) and (iii). The best fitting model was model (iv), the heritability, reduced model (bolded). The full model including genetic effects was an equally good fit (see Table S8 for full model results).

| Model | Fixed Effects | Random Effects | DIC | $\Delta$ DIC | $h^2$<br>[95% CI] | Residual Variance<br>[95% CI] |
| --- | --- | --- | --- | --- | --- | --- |
| (i) Heritability, Intercept Only | Intercept Only | a + m + g + c | 610.589 | 96.985 | 0.047<br>[0.002 - 0.122] | 0.773<br>[0.659 - 0.872] |
| (ii) Maternal Effects, Reduced | s + gs + oe | m | 578.858 | 65.254 | - | 0.979<br>[0.946, 0.998] |
| (iii) Environmental Effects, Reduced | s + gs + oe | m + g + c | 519.648 | 6.044 | - | 0.846<br>[0.765, 0.928] |
| **(iv) Heritability, Reduced** | **s + gs + oe** | **a + m + g + c** | **513.604** | 0.000 | **0.043**<br>**[0.002 - 0.110]** | **0.801**<br>**[0.683, 0.897]** |

## Discussion

We report three major findings about the development of affiliative social behavior in wild baboons. First, males and females differ in the age at which they first groom a conspecific, in their grooming relationships with their mothers, and in the environmental predictors of age at first grooming. Second, our results emphasize the important role of mothers in the development of social behavior: mothers were chosen as first grooming partners 19-fold more than expected by chance and were drivers of grooming relationships early in their offspring lives. Finally, age at first grooming is modestly but detectably heritable, suggesting that evolution by natural selection could shape this important developmental milestone (30).

### Sex differences in the development of grooming and grooming reciprocity

Females tended to groom conspecifics for the first time about two months earlier than males (Figure S1). Females also reached reciprocity with mothers at an earlier age than males, consistent with the idea that the early development of affiliative social behaviors and strong bonds with mothers may be more important for the philopatric sex (22, 24, 26, 66–70). Two of our results indicate that these sex differences in the development of social behavior are driven primarily by offspring behavior. First, daughters reached reciprocity with mothers well before sons did, even though mothers groomed sons and daughters at similar frequencies. Second, daughters directed grooming towards their mothers at much higher relative frequencies than sons. Previous work in other species has also found sex-differences in mother-offspring grooming patterns that are largely driven by daughters and not mothers (22, 24, 26, 66–69). Interestingly, the number of daughter-initiated grooming events peaks around menarche and then declines into adulthood. This decline may occur both because daughters increase their investment in relationships with other adult females (71) and because they begin to invest in their own offspring (median age of first birth is 5.97 years in the study population; 59).

### Environmental predictors of infant grooming patterns

Both males and females in larger social groups tended to begin grooming at later ages than those in small social groups. This could occur if adults in larger groups tend to engage in less affiliative behavior and more agonistic behavior – as suggested for yellow-bellied marmots -- potentially giving juveniles fewer opportunities to engage in grooming behavior (72). An alternative hypothesis is that larger groups contain more close-in-age juveniles who provide opportunities for non-grooming social interactions, such as play. This possibility is supported by a larger effect of group size on age at first grooming in males, as males tend to play more than females in many mammals, including primates (22, 73–75). Because sex differences in play behavior appear early in life, they likely lead to sex differences in the development of other social behaviors as well. Group size can also affect other aspects of a primate infant’s behavior that may explain differences in age of first grooming, including the proportion of time it spends in proximity to its mother and to other group members and the strength of social ties to peers (76, 77).

Beyond the consistent effect of group size in all our top models and in both sexes, we found modest evidence for other environmental effects on age at first grooming. For instance, maternal rank predicted male, but not female, age at first grooming (Table S2, Figure S4). We also found limited evidence that drought in the first year of life slightly accelerated age at first grooming for females (not males), but only after the median age at first grooming (Figure S3).

Although the presence of a sibling was not a significant predictor of the age at which a subject first groomed, the presence of kin clearly shaped subjects’ grooming patterns: 59% of first grooming partners were kin with r≥0.0625, and most of these (48% of the total) were male or female close kin (r<.25; Figures 1, S6). Mothers were chosen as first grooming partners much more than expected by chance, by both male and female subjects. This result is unsurprising, as young baboons spend the majority of their time with their mothers and mothers are key contributors to their offspring social development (78, 79). Together, our results are consistent with the extensively documented importance of mothers in particular, and kin in general, in affiliative relationships of primates, especially cercopithecines (80–82).

### Heritability of age at first grooming

Age at first grooming was weakly heritable. Thus, this trait is visible to natural selection, although the response to selection might be slow (30). In general, the heritability of behavioral traits is lower than that of morphological traits (36, 83). However, our value of h^2^=0.043 is much lower than the average of h^2^~0.30 reported in two meta-analyses of the heritability of social behavior (35, 84), and 4-6 times lower than the heritability of adult grooming behavior (h^2^=0.16-0.26) measured in our study population (41). Our low heritability estimate may partly reflect measurement error. First grooming events are relatively brief, one-time events in an individual’s life, which often must occur when observers are not with the group. In spite of the care with which our dataset was constructed, it is unlikely that we captured all first grooming events with high accuracy: as noted in the Methods, our measure of age at first grooming should be considered the latest date by which this milestone was achieved, rather than the exact date on which it was achieved. Thus, our non-zero heritability estimate is likely a lower bound on the true heritability for this behavioral milestone.

In our best fitting model, more than 75% of the variance in age at first grooming was unexplained. This unexplained variance is likely a consequence of three contributing factors. First, the trait we measured is likely to be an integrated trait, i.e., one that emerges from the combination of a large number of component traits (84, 85). The variation in each component of an integrated trait – which, for a social developmental trait, may include multiple traits involved in neurological and physical development – contributes to compounded residual variation in the integrated trait (84–88). Such compounded variation can be large, leading not only to large unexplained variation but also to low heritability, even if the trait has relatively high additive genetic variance (84, 86–88). Second, the large amount of unexplained variation in our model also points to unmeasured environmental effects, such as social proximity patterns, social network density, and mothering style (22, 23, 89), which represent potential future targets of study. Finally, it is likely that some of the unexplained variation in our model is attributable to indirect genetic effects, (IGEs) in which the genotypes of social partners affect focal phenotypes (Moore et al. 1997). Given the demonstrated importance of IGEs in other studies of behavior (41, 90, 91), the role of IGEs in social interactions in the wild represents another important topic for future research. In sum, our work provides valuable fundamental information on the development of primate social traits and their ability to evolve.

## Acknowledgements

In Kenya, our research was approved by the Kenya Wildlife Service, the Wildlife Research & Training Institute, the National Environment Management Authority, and the National Council for Science, Technology, and Innovation. We also thank the University of Nairobi, the Institute of Primate Research, the National Museums of Kenya, the members of the Amboseli-Longido pastoralist communities, the Enduimet Wildlife Management Area, Ker & Downey Safaris, Air Kenya, and Safarilink for their cooperation and assistance in the field. Particular thanks go to the Amboseli Baboon Research Project field team (R.S. Mututua, S. Sayialel, J.K. Warutere, I.L. Siodi, G. Marinka, B. Oyath) and camp staff. We also thank T. Wango and V. Oudu for their untiring assistance in Nairobi, and Jeanne Altmann for her fundamental contributions to the Amboseli baboon research. The baboon project database, Babase, was designed and programmed by K. Pinc and is expertly managed by N.H. Learn and J.B. Gordon. For a complete set of acknowledgments of funding sources, logistical assistance, and data collection and management, please visit http://amboselibaboons.nd.edu/acknowledgements/.

## Data Accessibility

All data used in this study will be archived on Duke University Digital Repository.

