## Supplemental Materials for "Heritable and sex-specific variation in the development of social behavior in a wild primate"

for

### Supplementary Methods and Analyses

*Description of predictors used in our model selection analysis*

Subject's sex: Infant sex was assigned within a few days of birth, based on visual inspection with binoculars from approximately 5 meters distance. Baboon genitalia are conspicuous, even in infants, and errors in the identification of sex are rare and quickly identified and corrected. No infant had an unidentified or incorrect sex for more than a few days after birth.

Maternal variables: Mothers are crucial to offspring development, so we included four maternal variables previously linked to mother-offspring social relationships or infant development in our models: maternal parity, maternal social isolation, and maternal dominance rank measured in two different ways (1, 2). Maternal parity was a binary variable indicating if the mother was primiparous (if the subject was its mother's first offspring) or multiparous (if the mother had previous offspring). Maternal social isolation was a binary variable indicating if a mother's social connectedness to other females was in the lowest quartile during the subject's first two years of life (3). We used two measures of maternal dominance rank (alpha status and proportional dominance rank) because these different dominance rank measures have been shown to have independent effects on traits in this population (4). Alpha status was a binary variable indicating whether or not the subject's mother was the top-ranking female in the group at any point in the period between birth and first observed grooming. Proportional dominance rank was measured continuously as the mean proportion of adult females the mother outranked from birth to the first time the subject was observed to groom another individual, with 1 corresponding to a top-ranking female who was ranked above all other adult females, and 0 corresponding to the lowest ranking female who was ranked below all other adult females (see Levy, Gesquiere (4) for a more detailed description of how dominance ranks are assigned).

Physical Environment: The Amboseli ecosystem is characterized by a predictable five month-long "dry season" (June through October) and a seven month-long "wet season" in which rain is highly variable and unpredictable. Overlaid on these seasonal changes, Amboseli sometimes experiences drought years in which rain is much lower than average or fails entirely. In our models we included two effects of the physical environment: (i) whether the subject was born during the wet versus the dry season, and (ii) whether or not the subject experienced drought in the first year of life (<200 mm of rainfall following Tung, Archie (3)). Both of these environmental effects are related to resource availability, which is limited during the dry season and in drought years, and may negatively affect offspring development (5, 6).

<u>Social Environment:</u> We included two measures of the subject's social environment in our models. First, we measured group size as the average number of individuals in the social group during the period from the subject's birth until the first time it was observed to groom another individual. Second, we recorded whether the subject had at least one maternal sibling co-residing in its social group for any amount of time between its birth and the first time it groomed another animal (mean 83.4% of time; range 10.5%-100%). We included this variable because maternal siblings are important grooming partners in adulthood (7, 8).

*Effects of hybridity on age at first grooming behavior*

The Amboseli baboons are a hybrid population of primarily yellow baboon (*Papio cynocephalus*) ancestry with genetic admixture from neighboring anubis baboon (*P. anubis*) populations. Individuals vary in their amount of hybrid ancestry and this variation is associated with developmental and social behavioral traits (9-13). Our most accurate measure of hybrid ancestry is based on genome-wide ancestry calls obtained from whole-genome resequencing data (12), but restricts data sets that include genetic hybridity to a subset of animals in the Amboseli population. Therefore, we first carried out the main analysis with the full data set, which included all the variables described above but not genetic hybrid score. We then conducted two secondary model selection analyses that included genetic hybrid scores.

Our genome-wide estimates of hybridity, generated in Vilgalys, Fogel (12), are measured as the proportion of an individual's genome derived from *P. anubis* ancestry, with a score of 1 corresponding to a fully *P. anubis* genome and a score of 0 corresponding to a fully *P. cynocephalus* genome (12). In brief, to estimate genome-wide ancestry, local ancestry was called along the genome using a composite likelihood method suitable for low coverage data (LCLAE: 14), using allele frequencies for unadmixed yellow and anubis baboons as a reference (15; see 12). Local ancestry calls were then averaged across the autosomes for each individual to produce a global, genome-wide estimate of anubis ancestry.

To understand the role of genetic hybridity on age at first grooming behavior, we ran two secondary model selection analyses that included the hybrid score of the subject and the hybrid score of the subject's mother, respectively, in addition to other environmental fixed effects. As noted in the main text and above, these datasets are smaller than the full dataset, because we have genetic hybrid scores for only a subset of subjects. Specifically, for our analysis of mothers' hybrid scores (N=148 mothers), we had hybrid scores for mothers of N=528 subjects, representing 250 males and 278 female subjects (mean hybrid score of mothers: 0.36 ± 0.09 SD; range: 0.24-0.60). For our analysis of subjects' hybrid scores, we had hybrid scores for N=237 subjects, representing 101 males and 136 females (mean hybrid score of subjects: 0.38 ± 0.09 SD; range: 0.23-0.71). In both of these analyses we retained all of the other fixed and random effects that were in the main analysis.

Similar to our models without hybrid scores, group size and observer effort were included in all top models. We also found that the subject's hybrid score was present in the best fitting models of age at first grooming (Table S3-S4), but maternal hybrid score was not (Tables S4-

S5). For both sexes, approximately half of the best fitting models included the subject's own hybrid score (females: 12 of the 22; males: 6 of 13), while maternal hybrid score was present only in one top model for females.

Although our model selection analyses lent support to the idea that individuals with more anubis ancestry tended to groom earlier, the model-averaged parameters indicated that the confidence intervals for the hazard ratio are very large and overlap one for both sexes (females: HR: 7.648, 95% CI: 0.680-85.983; males: HR: 7.716, 95% CI: 0.633-95.097; Table S4), indicating uncertainty as to whether this effect is biologically significant. The top model for females included, in addition to hybrid score, group size, observer effort, and presence of a sibling in the group; the top model for males included all of these effects plus maternal social isolation. However, the model-averaged hazard ratios for presence of a sibling and observer effort overlapped one for both males and females and overlapped one for the effect of maternal social isolation in males (Table S4). In aggregate, we conclude that these models provide strong support only for group size and observer effort as a fixed effect predicting age at first grooming. Nevertheless, whether the weak signal of genetic ancestry is robust in more highly powered analyses remains an intriguing question for future work, especially given previous findings indicating subtle, ancestry-linked differences in social behavior in the Amboseli population (9, 11, 13).

*Dispersal effects on male grooming reciprocity*

Our analyses of the development of grooming reciprocity are complicated by the fact that males vary in the age at which they disperse from the natal group and it is possible that males with stronger bonds with their mother delay dispersal. To confirm that our results were not dependent on male dispersal timing, we assessed if mother-offspring grooming relationships differed between males who dispersed before the median dispersal time (7.6 years) and those who dispersed after the median dispersal time. We find that there is no effect of dispersal time on reciprocity, number of grooming events initiated by the mother, the number of grooming events initiated by the offspring, or the total number of grooming events between each mother-offspring pair (Table S10, Figures S9, S10).

*Analysis of the genetic basis of age at first grooming including offspring hybrid score*

The main text describes an animal model to measure heritability of age at first grooming, using the full dataset with sex and environmental effects as fixed effects (see "Genetic variance in age at first grooming" in the Methods of the main text). We repeated this animal model to include hybrid score as a fixed effect, using the smaller subset of subjects for which hybrid score was available. In this dataset, the pedigree necessary to estimate heritability of age at first groom includes 485 individuals and a maximum of 5 generations, resulting in a trimmed pedigree with 323 individuals with known mothers, and 243 individuals with known fathers. In this trimmed pedigree we had 19 full sibling pairs, 241 maternal half sibling pairs, 316 paternal half sibling pairs, and 41 sibling pairs that were at least maternal half siblings but for whom the father of at least one of the members of the pair was unknown (i.e., some of these pairs could have been full siblings). Using the animal model approach described in the main text, our ‘full model’ with the trimmed pedigree included sex, social group size, observer effort, offspring hybrid score, presence of a sibling, and maternal social isolation by sex interaction. Our ‘reduced model’ with the trimmed pedigree included sex, social group size, observer effort, and offspring hybrid score. These models generate higher heritability estimates than the models excluding hybrid score, although the confidence intervals for $h^2$ overlap between models including and excluding hybrid score. Full results are presented in Table S9 and Figure S8.

**Supplementary Tables**

Table S1. Top models for environmental predictors of age at first observed grooming for female and male baboons. All N=781 individuals with an age at first observed grooming are included in this model. Models with Δ $AIC_c$<2 from the model with the lowest $AIC_c$ (“top models” discussed in the main text) are shown. Among the fixed effects, observer effort is denoted by “OE” and maternal variables are denoted by “M” with a subscript specifying the effect. See Tables S3 and S5 for results in a smaller, secondary analysis including hybrid scores.

| Sex | Fixed Effects | $AIC_c$ | Δ $AIC_c$ | Weight |
|---|---|---|---|---|
| (a) Female | Group Size + OE + Drought | 4098.799 | 0.000 | 0.070 |
| | Group Size + OE + Drought + $M_{Social\ Isolation}$ | 4099.367 | 0.568 | 0.053 |
| | Group Size + OE + Drought + Sibling | 4099.587 | 0.789 | 0.047 |
| | Group Size + OE + Drought + $M_{Parity}$ | 4099.879 | 1.081 | 0.041 |
| | Group Size + OE + Drought + Season | 4100.277 | 1.478 | 0.033 |
| | Group Size + OE + Drought + $M_{Prop\ Rank}$ | 4100.302 | 1.503 | 0.033 |
| (b) Male | Group Size + OE + $M_{Prop\ Rank}$ + $M_{Alpha}$ | 3586.545 | 0.000 | 0.084 |
| | Group Size + OE + $M_{Prop\ Rank}$ + $M_{Alpha}$ + Sibling | 3587.949 | 1.405 | 0.042 |
| | Group Size + OE | 3588.130 | 1.585 | 0.038 |
| | Group Size + OE + $M_{Prop\ Rank}$ + $M_{Alpha}$ + Drought | 3588.345 | 1.801 | 0.034 |
| | Group Size + OE + $M_{Prop\ Rank}$ + $M_{Alpha}$ + $M_{Parity}$ | 3588.358 | 1.814 | 0.034 |

Table S2. Model averaged parameter estimates for fixed effects that predict variance in age at first grooming. Estimates refer to change in the log hazard ratio and are calculated from the full coefficient set, but with terms not included in the top models set to zero. See Tables S4 and S6 for results in a smaller, secondary analysis including hybrid scores. Effects where 95% confidence intervals of the hazard ratio do not overlap one are bolded.

| Sex | Effect | Estimate | SE | HR | 95% CI HR | Interpretation |
|---|---|---|---|---|---|---|
| a) Female | **Group Size** | **-0.011** | **0.004** | **0.989** | **0.982 - 0.996** | ↑ group size = later age first grooming |
| | **Observer Effort** | **0.161** | **0.034** | **1.175** | **1.099 - 1.256** | ↑ observer effort = earlier age at first grooming |
| | Sibling Present | 0.016 | 0.060 | 1.017 | 0.879 - 1.379 | |
| | Maternal Social Isolation | -0.021 | 0.068 | 0.979 | 0.709 - 1.131 | |
| | Maternal Proportional Rank | 0.018 | 0.075 | 1.018 | 0.841 - 1.603 | |
| | Maternal Parity (primiparous) | -0.012 | 0.059 | 0.988 | 0.710 - 1.196 | |
| | **Drought in First Year of Life** | **0.404** | **0.186** | **1.498** | **1.041 - 2.155** | Experienced drought = earlier age at first grooming |
| | Season of Birth (wet) | -0.002 | 0.037 | 0.998 | 0.799 - 1.209 | |
| b) Male | **Group Size** | **-0.015** | **0.005** | **0.985** | **0.975 - 0.995** | ↑ group size = later age first grooming |
| | **Observer Effort** | **0.124** | **0.038** | **1.132** | **1.049 - 1.220** | ↑ observer effort = earlier age at first grooming |
| | Sibling Present | 0.021 | 0.069 | 1.021 | 0.881 - 1.432 | |
| | **Maternal Alpha Rank** | **0.387** | **0.259** | **1.473** | **1.048 - 2.410** | Alpha mom = earlier age at first grooming |
| | **Maternal Proportional Rank** | **-0.380** | **0.259** | **0.684** | **0.416 - 0.970** | ↑ Mom proportional rank = later age first grooming |
| | Maternal Parity (primiparous) | -0.014 | 0.067 | 0.986 | 0.675 - 1.216 | |
| | Drought in First Year of Life | -0.010 | 0.085 | 0.990 | 0.614 - 1.412 | |

Table S3. Top models for environmental predictors of age at first observed grooming for female and male baboons, including genetic hybrid score of the subject. N=231 individuals with an age at first observed grooming event had a genetic hybrid score are included in this model. Models with Δ $AIC_c$<2 from the top fitting model are shown. Maternal variables are denoted by "M". While genetic hybrid score appeared in several of the top models, the confidence intervals for its hazard ratio overlapped 1, indicating at most an effect too small to confirm in this data set (see Table S4).

| Sex | Fixed Effects | $AIC_c$ | Δ $AIC_c$ | Weight |
|---|---|---|---|---|
| a) Female | Group Size + OE + Hybrid Score + Sibling | 1057.969 | 0.000 | 0.024 |
| | Group Size + OE | 1057.982 | 0.014 | 0.024 |
| | Group Size + OE + Sibling | 1058.107 | 0.138 | 0.022 |
| | Group Size + OE + Hybrid Score + Sibling + Season | 1058.205 | 0.236 | 0.021 |
| | Group Size + OE + $M_{Parity}$ | 1058.552 | 0.583 | 0.018 |
| | Group Size + OE + Sibling + Drought | 1058.685 | 0.716 | 0.017 |
| | Group Size + OE + Hybrid Score | 1058.773 | 0.805 | 0.016 |
| | Group Size + OE + Sibling + Season | 1058.831 | 0.862 | 0.015 |
| | Group Size + OE + Hybrid Score + Sibling + $M_{Alpha}$ | 1058.887 | 0.918 | 0.015 |
| | Group Size + OE + Hybrid Score + Sibling + $M_{Alpha}$ + Season | 1058.988 | 1.019 | 0.014 |
| | Group Size + OE + Season | 1059.077 | 1.109 | 0.014 |
| | Group Size + OE + $M_{Parity}$ + Season | 1059.178 | 1.209 | 0.013 |
| | Group Size + OE + Hybrid Score + Sibling + $M_{Alpha}$ + $M_{Prop\ Rank}$ + Season | 1059.323 | 1.354 | 0.012 |
| | Group Size + OE + Hybrid Score + Sibling + $M_{Alpha}$ + $M_{Prop\ Rank}$ | 1059.475 | 1.506 | 0.011 |
| | Group Size + OE + Hybrid Score + Sibling + Drought | 1059.556 | 1.587 | 0.011 |
| | Group Size + OE + Sibling + $M_{Alpha}$ | 1059.720 | 1.751 | 0.010 |
| | Group Size + OE + $M_{Social\ Isolation}$ | 1059.722 | 1.753 | 0.010 |
| | Group Size + OE + Hybrid Score + Season | 1059.789 | 1.820 | 0.010 |
| | Group Size + OE + $M_{Alpha}$ | 1059.822 | 1.853 | 0.009 |
| | Group Size + OE + Hybrid Score + $M_{Parity}$ | 1059.881 | 1.913 | 0.009 |
| | Group Size + OE + Hybrid Score + Sibling + $M_{Prop\ Rank}$ | 1059.908 | 1.940 | 0.009 |
| | Group Size + OE + Hybrid Score + Sibling + $M_{Social\ Isolation}$ | 1059.927 | 1.958 | 0.009 |
| b) Male | Group Size + OE + Hybrid Score + Sibling + $M_{Social\ Isolation}$ | 715.758 | 0.000 | 0.038 |
| | Group Size + OE + Sibling + $M_{Social\ Isolation}$ | 716.038 | 0.281 | 0.033 |
| | Group Size + OE + $M_{Parity}$ + $M_{Social\ Isolation}$ | 716.624 | 0.867 | 0.025 |
| | Group Size + OE + Sibling | 716.768 | 1.011 | 0.023 |
| | Group Size + OE + Hybrid Score + Sibling | 716.778 | 1.020 | 0.023 |
| | Group Size + OE + Hybrid Score + $M_{Parity}$ + $M_{Social\ Isolation}$ | 717.083 | 1.326 | 0.020 |
| | Group Size + OE + Hybrid Score + $M_{Social\ Isolation}$ | 717.271 | 1.513 | 0.018 |
| | Group Size + OE | 717.307 | 1.550 | 0.018 |
| | Group Size + OE + $M_{Social\ Isolation}$ | 717.337 | 1.579 | 0.017 |
| | Group Size + OE + $M_{Parity}$ | 717.341 | 1.583 | 0.017 |
| | Group Size + OE + Hybrid Score | 717.364 | 1.606 | 0.017 |
| | Group Size + OE + Sibling + $M_{Social\ Isolation}$ + $M_{Alpha}$ | 717.646 | 1.888 | 0.015 |
| | Group Size + OE + Hybrid Score + $M_{Parity}$ | 717.754 | 1.996 | 0.014 |

Table S4. Model averaged parameter estimates for fixed effects that predict age at first grooming event in models that include the subject's genetic hybrid score. Estimates refer to change in the log hazard ratio and are calculated from the full coefficient set with terms set to zero when not included in the model. Note that, for subject's genetic hybrid score, the confidence intervals for the hazard ratio overlap 1, indicating at most an effect too small to confirm in this data set. Effects where 95% confidence intervals of the hazard ratio do not overlap one are bolded.

| Sex | Effect | Estimate | SE | HR | 95% CI HR | Interpretation |
|---|---|---|---|---|---|---|
| a) Female | **Group Size** | **-0.018** | **0.007** | **0.982** | **0.969-0.995** | ↑ group size = later age first grooming |
| | **Observer Effort** | **0.097** | **0.042** | **1.102** | **1.015-1.197** | ↑ observer effort = earlier age at first grooming |
| | Sibling Present | 0.273 | 0.283 | 1.314 | 0.996-2.461 | |
| | Maternal Social Isolation | -0.009 | 0.063 | 0.991 | 0.571-1.307 | |
| | Maternal Alpha Rank | -0.122 | 0.316 | 0.885 | 0.235-1.478 | |
| | Maternal Proportional Rank | 0.057 | 0.203 | 1.059 | 0.864-3.480 | |
| | Maternal Parity (primiparous) | -0.030 | 0.121 | 0.97 | 0.476-1.306 | |
| | Subject's Genetic Hybrid Score | 1.048 | 1.349 | 2.851 | 0.680-85.983 | |
| | **Drought in First Year of Life** | **0.080** | **0.279** | **1.083** | **1.207-5.105** | Experienced drought = earlier age at first grooming |
| | Season of Birth (wet) | -0.069 | 0.147 | 0.933 | 0.556-1.166 | |
| b) Male | **Group Size** | **-0.024** | **0.009** | **0.976** | **0.959-0.994** | ↑ group size = later age first grooming |
| | **Observer Effort** | **0.172** | **0.058** | **1.187** | **1.060-1.330** | ↑ observer effort = earlier age at first grooming |
| | Sibling Present | 0.230 | 0.298 | 1.259 | 0.986-2.667 | |
| | Maternal Social Isolation | 0.313 | 0.337 | 1.367 | 0.972-2.932 | |
| | Maternal Alpha Rank | 0.009 | 0.093 | 1.009 | 0.575-2.426 | |
| | Maternal Parity (primiparous) | -0.123 | 0.252 | 0.884 | 0.360-1.124 | |
| | Subject's Genetic Hybrid Score | 0.955 | 1.342 | 2.598 | 0.633-94.097 | |

Table S5. Top models for environmental predictors of age at first observed grooming for female and male baboons, including genetic hybrid score of the subject's mother. N=528 individuals with an age at first observed grooming event and whose mother had a genetic hybrid score are included in this model. Models with $\Delta$ $AIC_c$<2 from the top fitting model are shown. Maternal variables are denoted by "M". Maternal genetic hybrid score appears in only one of the top models for females, and none for males.

| Sex | Fixed Effects | $AIC_c$ | $\Delta$ $AIC_c$ | Weight |
|---|---|---|---|---|
| a) Female | Group Size + OE | -1256.804 | 2532.854 | 0.000 |
| | Group Size + OE + Sibling | -1255.790 | 2533.175 | 0.321 |
| | Group Size + OE + $M_{Parity}$ | -1256.312 | 2533.342 | 0.488 |
| | Group Size + OE + $M_{Hybrid\ Score}$ | -1256.973 | 2534.483 | 1.629 |
| | Group Size + OE + Season | -1256.464 | 2534.598 | 1.745 |
| | Group Size + OE + $M_{Social\ Isolation}$ | -1256.796 | 2534.700 | 1.846 |
| b) Male | OE + $M_{Parity}$ | 2240.350 | 0.000 | 0.024 |
| | OE + $M_{Parity}$ + Drought | 2240.466 | 0.116 | 0.022 |
| | OE + Group Size + $M_{Parity}$ | 2241.105 | 0.755 | 0.016 |
| | OE + $M_{Parity}$ + $M_{Alpha}$ | 2241.170 | 0.820 | 0.016 |
| | OE | 2241.188 | 0.838 | 0.015 |
| | OE + $M_{Parity}$ + $M_{Alpha}$ + Drought | 2241.345 | 0.995 | 0.014 |
| | OE + $M_{Alpha}$ | 2241.737 | 1.388 | 0.012 |
| | OE + $M_{Parity}$ + $M_{Social\ Isolation}$ | 2241.778 | 1.428 | 0.012 |
| | OE + Drought | 2241.865 | 1.515 | 0.011 |
| | OE + $M_{Parity}$ + $M_{Social\ Isolation}$ + Drought | 2241.907 | 1.557 | 0.011 |
| | OE + Group Size + $M_{Parity}$ + Drought | 2241.952 | 1.602 | 0.011 |
| | OE + $M_{Parity}$ + Sibling | 2242.000 | 1.650 | 0.010 |
| | OE + Group Size | 2242.033 | 1.683 | 0.010 |
| | OE + Group Size + $M_{Parity}$ + $M_{Alpha}$ | 2242.051 | 1.701 | 0.010 |

Table S6. Model averaged parameter estimates for fixed effects that predict age at first grooming event in models that include genetic hybrid score of the subject’s mother. N=528 individuals with an age at first observed grooming event and whose mother had a genetic hybrid score are included. Estimates refer to change in the log hazard ratio and are calculated from the full coefficient set with terms set to zero when not included in the model. Note that the estimate of the hazard ratio for the effect of maternal hybrid score is near one for females; it does not appear in any top models for males (Table S5). Effects where 95% confidence intervals of the hazard ratio do not overlap one are bolded.

| Sex | Effect | Estimate | SE | HR | 95% CI HR | Interpretation |
|---|---|---|---|---|---|---|
| a) Female | Group Size | -0.008 | 0.004 | 0.992 | 0.984-1.001 | |
| | **Observer Effort** | **0.227** | **0.036** | **1.255** | **1.171-1.346** | ↑ observer effort = earlier age at first grooming |
| | Sibling Present | 0.041 | 0.101 | 1.042 | 0.915-1.587 | |
| | Maternal Social Isolation | -0.004 | 0.047 | 0.996 | 0.727-1.276 | |
| | Maternal Parity (primiparous) | -0.038 | 0.106 | 0.963 | 0.597-1.148 | |
| | Maternal Genomic Hybrid Score | -0.037 | 0.287 | 0.963 | 0.152-3.412 | |
| | Season of Birth (wet) | -0.009 | 0.049 | 0.991 | 0.719-1.186 | |
| b) Male | Group Size | -0.002 | 0.004 | 0.998 | 0.981-1.002 | |
| | **Observer Effort** | **0.240** | **0.050** | **1.271** | **1.151-1.403** | ↑ observer effort = earlier age at first grooming |
| | Sibling Present | -0.009 | 0.062 | 0.991 | 0.562-1.246 | |
| | Maternal Social Isolation | -0.009 | 0.060 | 0.991 | 0.675-1.273 | |
| | Maternal Alpha Rank | 0.083 | 0.196 | 1.086 | 0.803-2.315 | |
| | Maternal Parity (primiparous) | -0.226 | 0.218 | 0.798 | 0.498-1.100 | |
| | Drought in First Year of Life | -0.079 | 0.171 | 0.924 | 0.516-1.243 | |

Table S7. Effects from generalized linear mixed models for number of grooms initiated by mothers versus offspring. Age (measured in months) is centered and standardized before analyses so estimates represent a change in number of grooms for a one SD increase in age.

| | | 0 - 4.5 years | | | | 4.5 - 7.7 years | | | |
|---|---|---|---|---|---|---|---|---|---|
| Grooming Measure | Effect | estimate | SE | Z value | P value | estimate | SE | Z value | P value |
| (A) Offspring Initiated | Sex (male) | -1.289 | 0.073 | -17.62 | <0.001 | -1.055 | 0.219 | -4.82 | <0.001 |
| | Age | 1.221 | 0.024 | 50.88 | <0.001 | -0.476 | 0.024 | -19.93 | <0.001 |
| | $Age^2$ | -0.759 | 0.019 | -40.06 | <0.001 | 0.114 | 0.025 | 4.64 | <0.001 |
| | Sex*Age | -0.305 | 0.042 | -7.29 | <0.001 | 0.125 | 0.045 | 2.80 | 0.005 |
| | Sex*$Age^2$ | 0.186 | 0.034 | 5.44 | <0.001 | -0.179 | 0.047 | -3.77 | <0.001 |
| (B) Mother Initiated | Sex (male) | -0.014 | 0.036 | -0.40 | 0.690 | -0.595 | 0.194 | -3.06 | 0.002 |
| | Age | -0.666 | 0.009 | -72.57 | <0.001 | -0.276 | 0.025 | -11.24 | <0.001 |
| | $Age^2$ | -0.097 | 0.009 | -10.32 | <0.001 | 0.131 | 0.026 | 5.08 | <0.001 |
| | Sex*Age | -0.148 | 0.013 | -11.02 | <0.001 | -0.201 | 0.044 | -4.57 | <0.001 |
| | Sex*$Age^2$ | -0.045 | 0.014 | -3.19 | 0.001 | -0.143 | 0.047 | -3.04 | 0.002 |
| (C) Total Grooms | Sex (male) | -0.265 | 0.039 | -6.89 | <0.001 | -0.777 | 0.233 | -3.33 | 0.001 |
| | Age | -0.346 | 0.009 | -39.27 | <0.001 | -0.441 | 0.024 | -18.42 | <0.001 |
| | $Age^2$ | -0.061 | 0.009 | -6.92 | <0.001 | 0.143 | 0.024 | 5.87 | <0.001 |
| | Sex*Age | -0.321 | 0.013 | -24.61 | <0.001 | -0.007 | 0.041 | -0.17 | 0.865 |
| | Sex*$Age^2$ | -0.022 | 0.014 | -1.64 | 0.100 | -0.159 | 0.043 | -3.69 | <0.001 |

Table S8. Comparison of results from nested animal models (full model) where fixed effects included sex (s), social group size (gs), observer effort (oe), presence of siblings (sib), drought x sex interaction (d*s), maternal alpha rank by sex interaction ($m_a$*s), and maternal proportional rank by sex interaction ($m_r$*s). In maternal effects models, only maternal ID (m) was included as a random effect. In early environmental effects models, random effects included maternal ID (m), social group ID (g), and cohort (c, based on hydrological year of birth). In heritability models, animal breeding value (a) was estimated as a random effect in addition to maternal ID, social group ID, and cohort. The best fitting model was the heritability model with reduced fixed effects (see Table 2), but the full heritability model had a Deviance Information Criteria (DIC) within 2 of the best model and so is bolded below. Heritability ($h^2$) and 95% Credible Interval (CI) for $h^2$ from the models that estimated it are also reported. See Table 2 for reduced and intercept only model results.

| Model | Fixed Effects | Random Effects | DIC | Δ DIC | $h^2$ [95% CI] | Residual Variance [95% CI] |
|---|---|---|---|---|---|---|
| Maternal Effects, Full | s + gs + oe + sib + d*s + $m_a$*s + $m_r$*s | m | 586.100 | 72.496 | - | 0.978 [0.945, 0.998] |
| Early Environmental Effects, Full | s + gs + oe + sib + d*s + $m_a$*s + $m_r$*s | m + g + c | 520.380 | 6.776 | - | 0.832 [0.739, 0.911] |
| **Heritability, Full** | **s + gs + oe + sib + d*s + $m_a$*s + $m_r$*s** | **a + m + g + c** | **513.869** | 0.265 | **0.042 [0.002 - 0.110]** | **0.784 [0.670, 0.888]** |

Table S9. Comparison of results from nested animal models from the analysis on a subset of subjects with hybrid scores. In the intercept only model we included only an intercept as a fixed effect. In reduced models, fixed effects included sex (s), social group size (gs), observer sampling effort (oe), presence of siblings (sib), and offspring hybrid score (as). In full models, fixed effects included sex, social group size, observer sampling effort, presence of siblings, offspring hybrid score, mother proportional rank ($m_r$), mother alpha rank status ($m_a$), and drought (d). In maternal effects models, only maternal ID (m) was included as a random effect. In early environmental effects models, random effects included maternal ID, social group ID, and cohort (c) measured by hydrological year. In heritability models, animal (a) was included as a random effect in addition to maternal ID, social group ID, and cohort as random effects. The best fitting model was the heritability with full fixed effects (bolded). Deviance Information Criteria (DIC) and ΔDIC are reported for all models. Heritability ($h^2$) and 95% Credible Interval (CI) for $h^2$ estimated from the three models that included animal as a random effect are also reported.

| Model | Fixed Effects | Random Effects | DIC | Δ DIC | $h^2$ [95% CI] | Residual Variance [95% CI] |
|---|---|---|---|---|---|---|
| Heritability, Intercept Only | Intercept Only | a + m + g + c | 163.286 | 44.910 | 0.121 [0.002 - 0.334] | 0.669 [0.408, 0.897] |
| Maternal Effects, Reduced | s + gs + oe + as | m | 153.795 | 35.419 | - | 0.966 [0.906, 0.998] |
| Early Environmental Effects, Reduced | s + gs + oe + as | m + g + c | 143.605 | 25.229 | - | 0.822 [0.662, 0.962] |
| Heritability, Reduced | s + gs + oe + as | a + m + g + c | 125.799 | 7.423 | 0.117 [0.002 - 0.301] | 0.677 [0.416, 0.919] |
| Maternal Effects, Full | s + gs + oe + as + sib + $m_{si}$*s | m | 150.164 | 31.788 | - | 0.962 [0.892, 0.998] |
| Early Environmental Effects, Full | s + gs + oe + as + sib + $m_{si}$*s | m + g + c | 138.877 | 20.501 | - | 0.817 [0.668, 0.953] |
| **Heritability, Full** | **s + gs + oe + as + sib + $m_{si}$*s** | **a + m + g + c** | **118.376** | **0** | **0.125 [0.001-0.307]** | **0.658 [0.381, 0.881]** |

Table S10. Analyses of mother-son reciprocity for early versus late dispersing males ("Dispersal (early)" effect).

| | | 0 - 4.5 years | | | | 4.5 - 7.7 years | | | |
|---|---|---|---|---|---|---|---|---|---|
| Grooming Measure | Effect | estimate | SE | Z value | P value | estimate | SE | Z value | P value |
| (a) Reciprocity | Dispersal (early) | 0.004 | 0.027 | 0.15 | 0.881 | -0.071 | 0.103 | -0.69 | 0.491 |
| | Age | -0.179 | 0.017 | -10.45 | <0.001 | -0.015 | 0.079 | -0.19 | 0.851 |
| | $Age^2$ | -0.042 | 0.017 | -2.46 | 0.014 | 0.036 | 0.082 | 0.45 | 0.655 |
| | Dispersal *Age | 0.007 | 0.022 | 0.32 | 0.752 | -0.049 | 0.088 | -0.56 | 0.579 |
| | Dispersal *$Age^2$ | 0.003 | 0.022 | 0.15 | 0.885 | 0.005 | 0.092 | 0.06 | 0.953 |
| (b) Offspring Initiated | Dispersal (early) | -0.041 | 0.124 | -0.33 | 0.742 | 0.420 | 0.286 | 1.47 | 0.143 |
| | Age | 0.969 | 0.060 | 16.24 | <0.001 | -0.478 | 0.089 | -5.34 | <0.001 |
| | $Age^2$ | -0.596 | 0.051 | -11.67 | <0.001 | -0.297 | 0.095 | -3.12 | 0.002 |
| | Dispersal *Age | -0.018 | 0.077 | -0.23 | 0.818 | 0.265 | 0.101 | 2.63 | 0.009 |
| | Dispersal *$Age^2$ | 0.055 | 0.066 | 0.83 | 0.409 | 0.290 | 0.107 | 2.72 | 0.007 |
| (c) Mother Initiated | Dispersal (early) | 0.013 | 0.067 | 0.20 | 0.843 | 0.265 | 0.253 | 1.05 | 0.296 |
| | Age | -0.754 | 0.019 | -38.90 | <0.001 | -0.538 | 0.080 | -6.73 | <0.001 |
| | $Age^2$ | -0.143 | 0.019 | -7.38 | <0.001 | -0.286 | 0.086 | -3.31 | 0.001 |
| | Dispersal *Age | 0.068 | 0.024 | 2.80 | 0.005 | 0.167 | 0.091 | 1.85 | 0.065 |
| | Dispersal *$Age^2$ | 0.032 | 0.025 | 1.26 | 0.207 | 0.358 | 0.097 | 3.68 | <0.001 |
| (d) Total Grooms | Dispersal (early) | 0.014 | 0.069 | 0.20 | 0.841 | 0.387 | 0.274 | 1.41 | 0.158 |
| | Age | -0.573 | 0.018 | -31.32 | <0.001 | -0.555 | 0.081 | -6.87 | <0.001 |
| | $Age^2$ | -0.087 | 0.019 | -4.65 | <0.001 | -0.311 | 0.087 | -3.57 | <0.001 |
| | Dispersal *Age | 0.050 | 0.023 | 2.15 | 0.032 | 0.240 | 0.091 | 2.62 | 0.009 |
| | Dispersal *$Age^2$ | 0.024 | 0.024 | 0.98 | 0.329 | 0.362 | 0.098 | 3.69 | <0.001 |

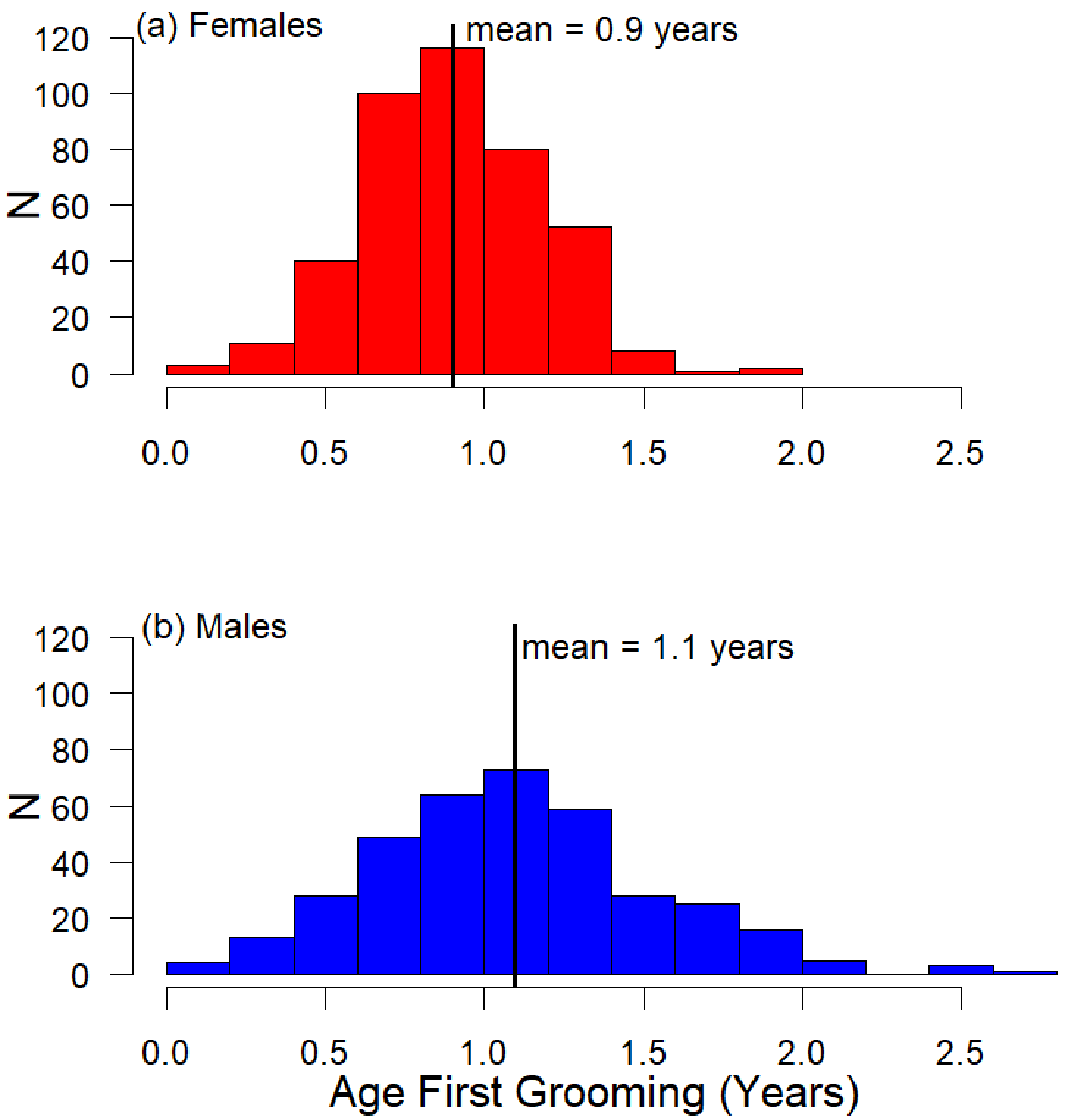


Figure S1. Distributions of age at first grooming for (a) female and (b) male subjects. Females groomed earlier than males (GLMM: b=0.189 ± 0.025, z=0.7599; p<0.001), with females first grooming at an average age of 0.9 years (95%CI: 0.83-0.96 years) and males first grooming at an average age of 1.1 years (95%CI: 1.02-1.15 years). Note that because grooming events are relatively brief, one-time events, which may occur when observers are not recording behavior, the actual first grooming behavior may be earlier than the age at first observed grooming we show here.

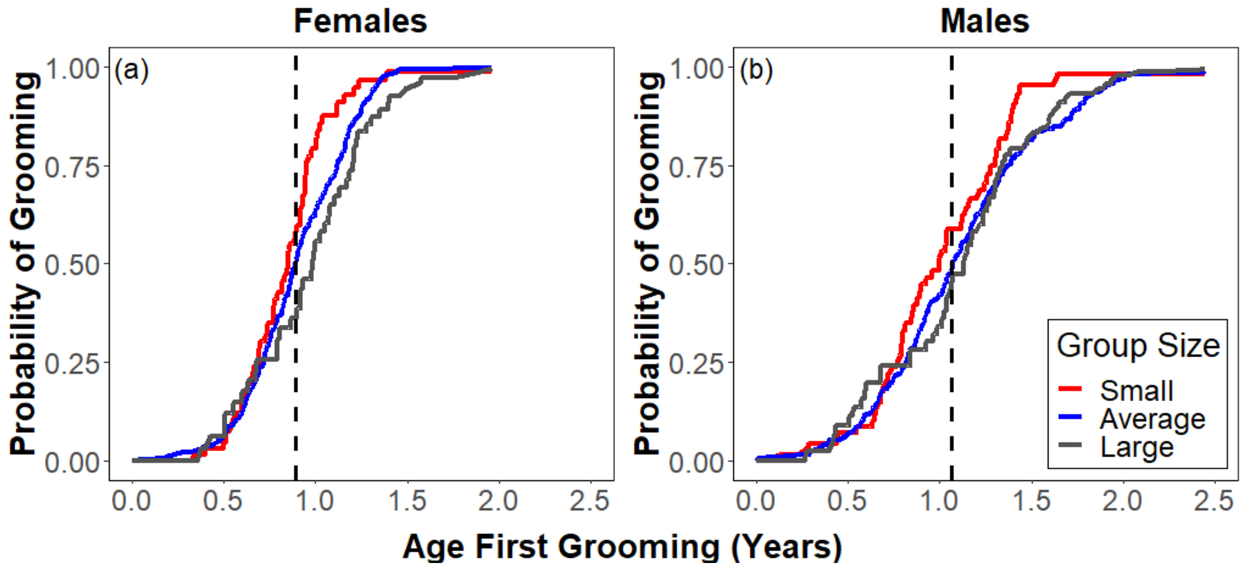


Figure S2. Group size predicts age at first grooming for both sexes, appearing in all of the top Cox proportional hazards models for both males and females. Both (a) females and (b) males in smaller groups groomed earlier than those in larger groups. Predicted values are plotted. Group size was treated as a continuous variable in our models; we stratified it into three categories for visualization purposes only, with small group size defined as groups more than 1 SD below the mean size (N=62 females, 55 males), average group size defined as groups within 1 SD of the mean (N=307 females, 254 males), and large group size defined as groups more than 1 SD above the mean group size (N=44 females, 59 males). The dashed vertical line depicts the median (and mean) age at first grooming (females: 0.9 years, males: 1.1 years).

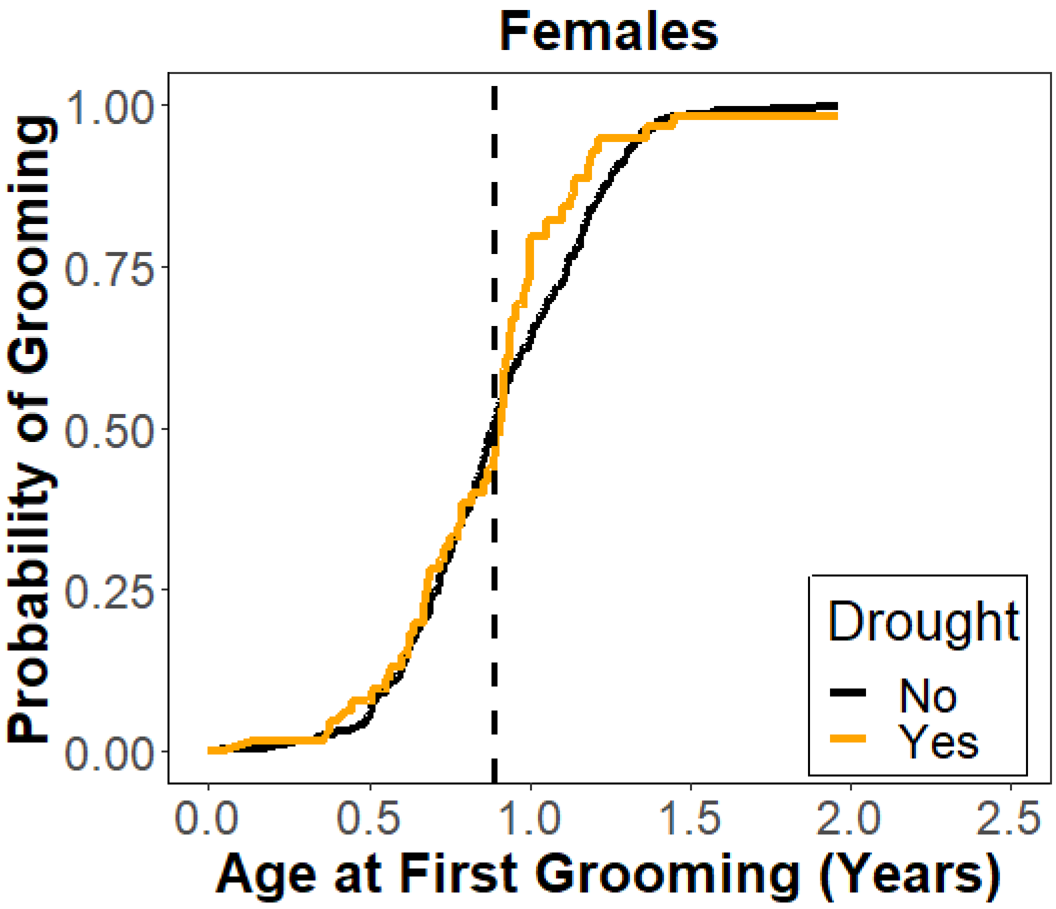


Figure S3. The effect of experiencing drought in the first year of life on age at first grooming event for females (N=53 experienced drought, 360 females did not experience drought). Raw values are plotted. Predicted values are plotted. The dashed vertical line depicts the median (and mean) age at first grooming for females (0.9 years).

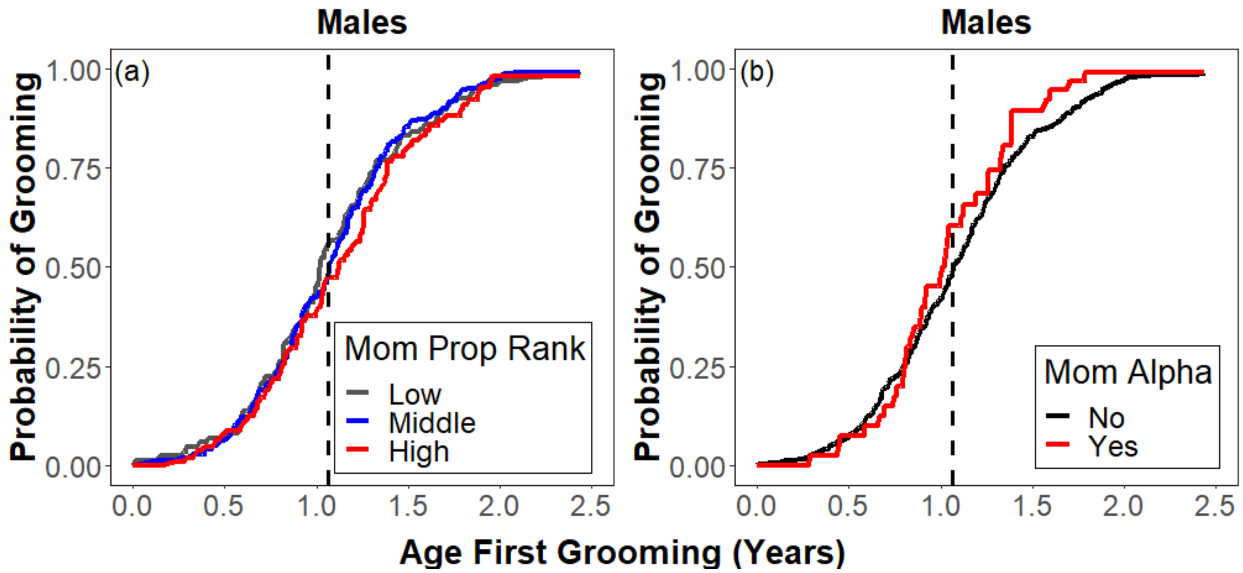


Figure S4. The effect of (a) maternal proportional rank and (b) maternal alpha status on male age at first grooming event (N=38 males with alpha mothers, 330 males with non-alpha mothers). Predicted values are plotted. Low maternal proportional rank corresponds to males with mothers whose rank was in the lowest quartile (N=93), middle maternal proportional rank corresponds to males with mothers whose rank was in the middle two quartiles (N=185), and high maternal proportional rank corresponds to males with mothers whose rank was in the highest quartile (N=90). The dashed vertical line depicts the median (and mean) age at first grooming for males (1.1 years).

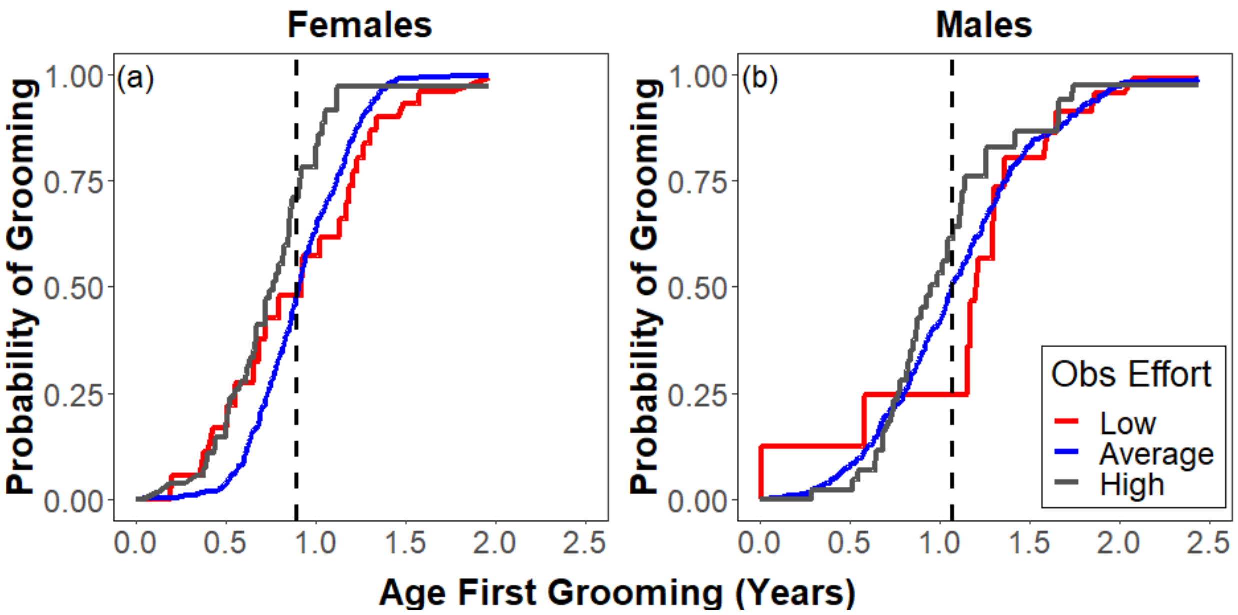


Figure S5. The effect of observer effort on age at first grooming event for (a) females and (b) males. Predicted values are plotted. Low observer effort was defined as groups with per capita grooming less than 1 SD below the mean (N=24 females, 12 males), average observer effort was defined as groups with per capita grooming within 1 SD of the mean (N=343 females, 320 males), and high observer effort were groups with per capita grooming more than 1 SD above the mean (N=46 females, 36 males). The dashed vertical line depicts the median (and mean) age at first grooming (females: 0.9 years, males: 1.1 years).

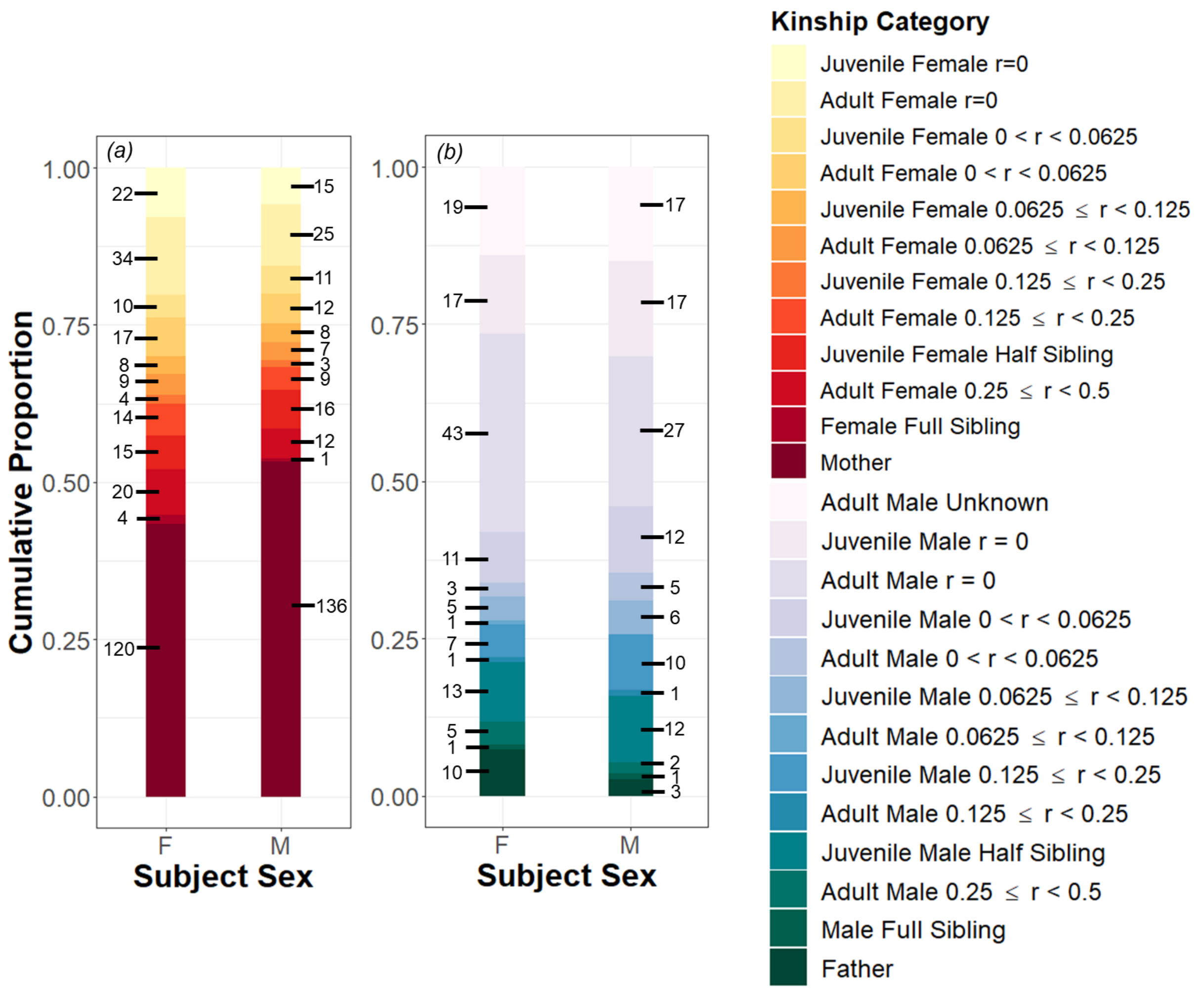


Figure S6. Finer scale breakdown of the identities of first grooming partners for female and male study subjects. Panel (a) shows first grooming partners who were females; females were the first observed grooming partners for 67% of female subjects (n=277) and 69% of male subjects (n=255). Panel (b) shows first grooming partners who were males; 33% of female subjects and 31% of male subjects first groomed a male. Numbers next to each bar denote the number of first grooming partners in that category. First grooming partners were categorized by sex, age, and pedigree-based kinship category. Age categories included (i) juvenile males (aged <7 years, category includes subadults), (ii) juvenile females (aged <4 years), (iii) adult males, and (iv) adult females.  Kinship categories included (i) r=0.5 (mothers, fathers, full siblings), (ii) 0.25 ≤ r 0.5 (e.g., half siblings, grandmothers, full aunts), (iii) 0.125 ≤ r < 0.25 (e.g., full cousins, great grandparents, half-aunts, half-uncles, half-nieces, half-nephews), (iv) 0.0625 ≤ r < 0.125 (e.g., half-cousins, great-great grandparents), 0 < r < 0.0625 (e.g., second cousins, half second cousins), unrelated (r = 0), and unknown (applies only to immigrant males with no known offspring or adult relative in the group). The number of fathers and paternal relatives identified as first grooming partners is underestimated, because many subjects (n=292) lacked paternity assignments.

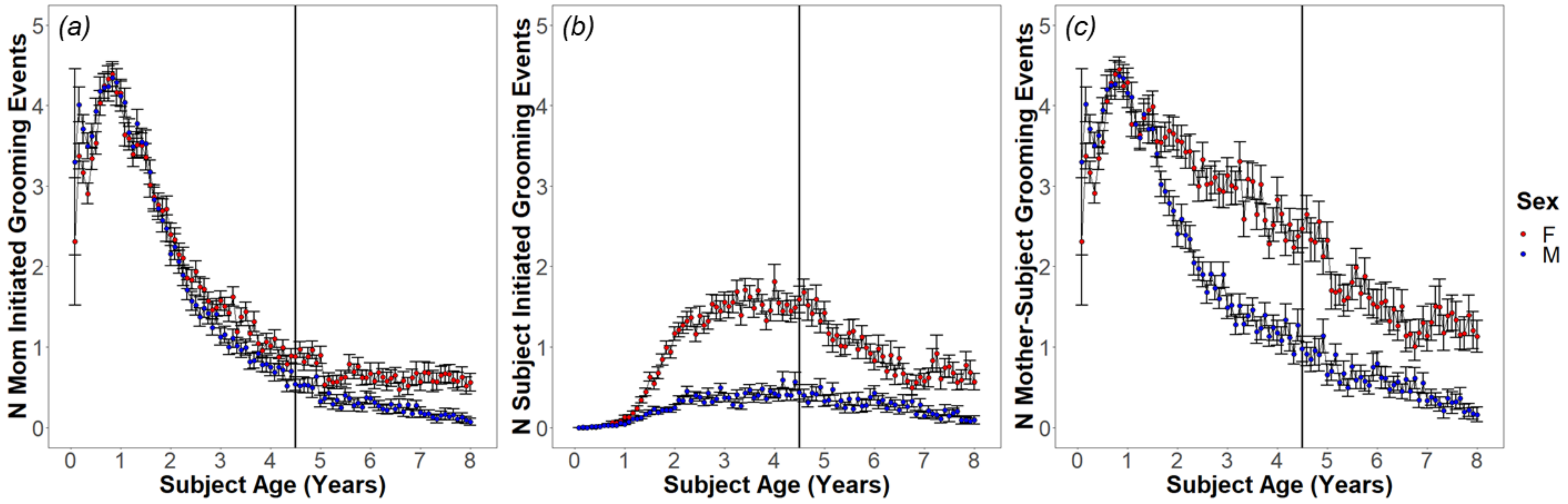


Figure S7. Number of mother-offspring grooming events initiated by the (a) mother to their daughter (red) or son (blue) or (b) initiated by daughter or son to their mother as a function of offspring age. The total number of mother-offspring grooming events is shown in panel (c). Points are monthly averages and bars denote standard error. The vertical black line represents the average age at female menarche (4.5 years).

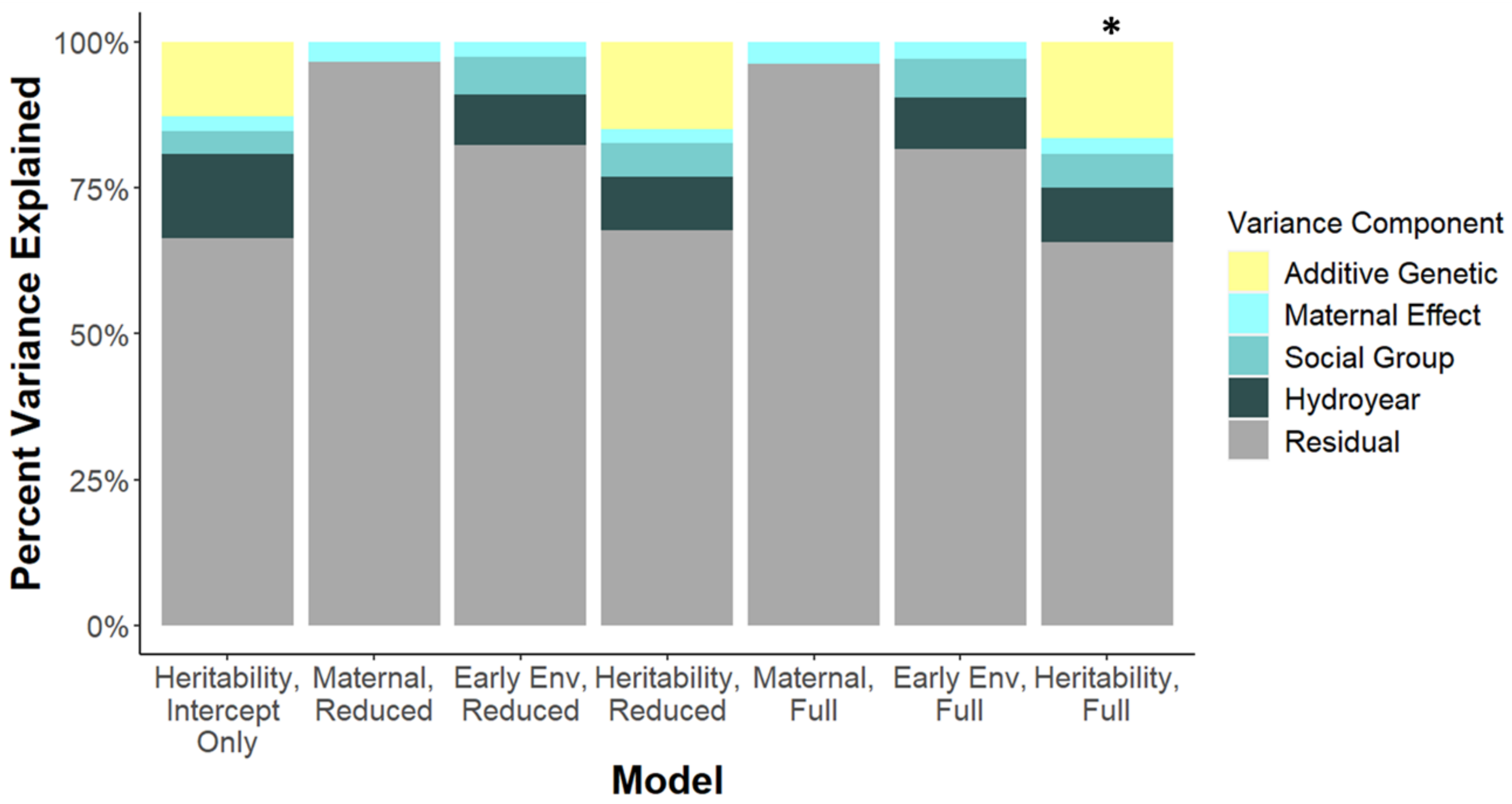


Figure S8. Percent of variance in age at first groom explained by additive genetic, maternal, social group, hydrological year, and residual effects in models that included the individual's hybrid score. The best fitting model (indicated by a star) included additive genetic effects, maternal effects, social group effects, and hydrological year effects. All models with additive genetic effects fit better than nested models without. See Table S9 and Supplementary Analyses: "*Analysis of the genetic basis of age at first grooming including offspring hybrid score*" text for definitions of models.

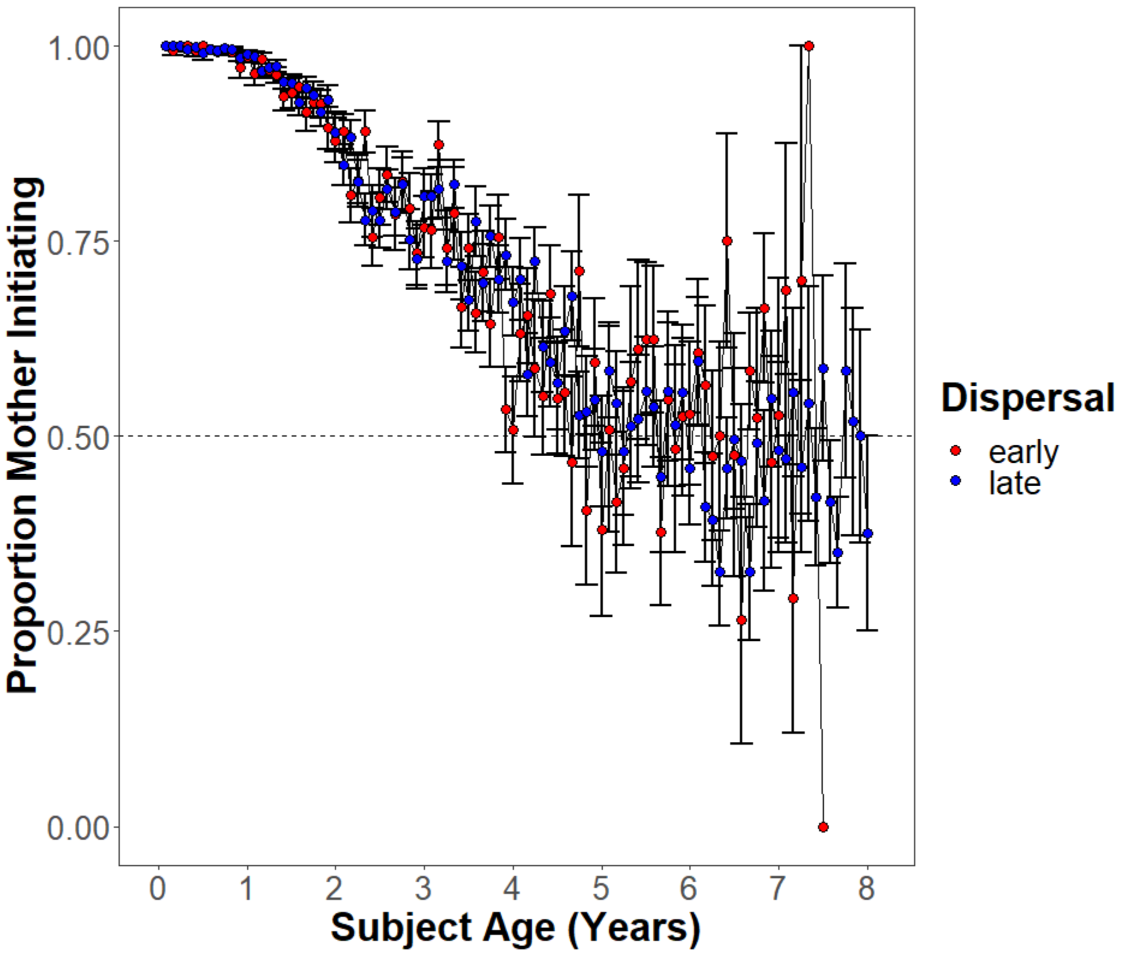


Figure S9. Proportion of mother-son grooming events where the mother-initiated grooming events as a function of age for males that dispersed early, before the median age at dispersal (7.6 years; red) and males that dispersed late, after the median age at dispersal (blue). The dashed line indicates perfect reciprocity where the number of grooms by the mother and son are equal. There is no effect of early versus late dispersal on mother-son reciprocity (Table S10).

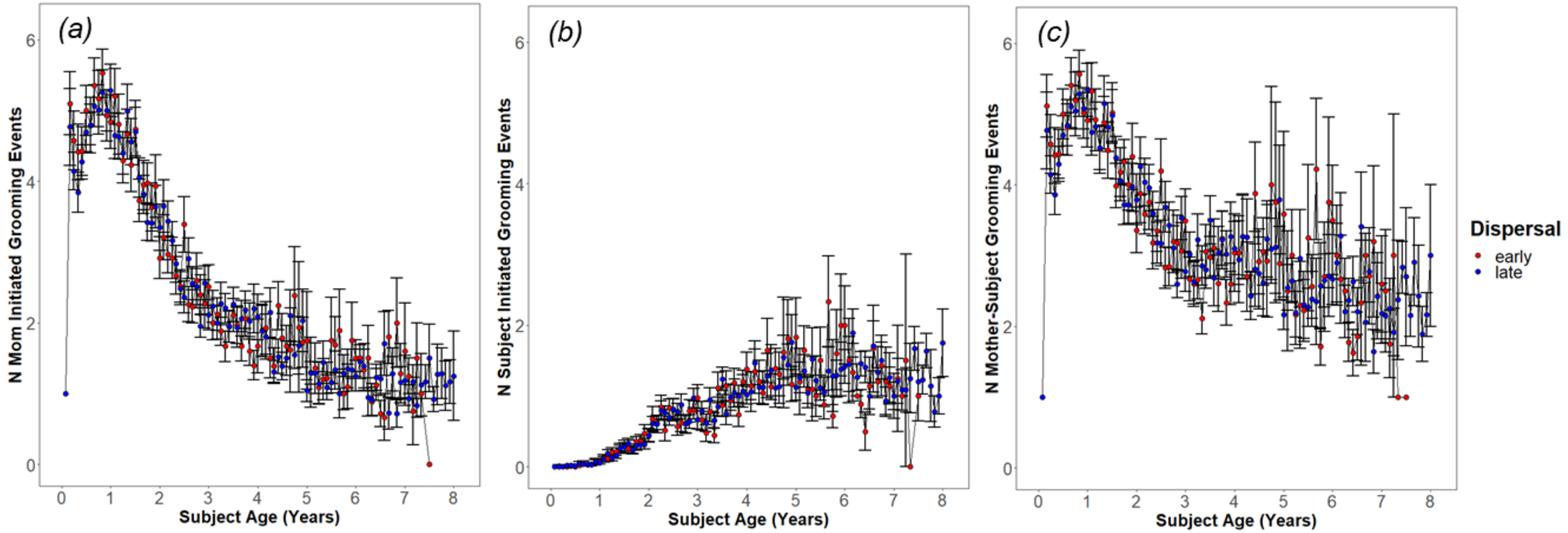


Figure S10. Number of mother-son grooming events initiated by the (a) mother or (b) son changes with offspring age. The total number of mother-offspring grooming events is shown in panel (c). Averages for males that dispersed early, before the median age at dispersal (7.6 years) are shown in red and males that dispersed late, after the median age at dispersal are shown in blue.
